# Assembly of complete diploid phased chromosomes from draft genome sequences

**DOI:** 10.1101/2021.11.11.468134

**Authors:** Andrea Minio, Noé Cochetel, Amanda Vondras, Mélanie Massonnet, Dario Cantu

## Abstract

*De novo* genome assembly is essential for genomic research. High-quality genomes assembled into phased pseudomolecules are challenging to produce and often contain assembly errors because of repeats, heterozygosity, or the chosen assembly strategy. Although algorithms that produce partially phased assemblies exist, haploid draft assemblies that may lack biological information remain favored because they are easier to generate and use. We developed HaploSync, a suite of tools that produces fully phased, chromosome-scale diploid genome assemblies, and performs extensive quality control to limit assembly artifacts. HaploSync scaffolds sequences from a draft diploid assembly into phased pseudomolecules guided by a genetic map and/or the genome of a closely related species. HaploSync generates a report that visualizes the relationships between current and legacy sequences, for both haplotypes, and displays their gene and marker content. This quality control helps the user identify misassemblies and guides Haplosync’s correction of scaffolding errors. Finally, HaploSync fills assembly gaps with unplaced sequences and resolves collapsed homozygous regions. In a series of plant, fungal, and animal kingdom case studies, we demonstrate that HaploSync efficiently increases the assembly contiguity of phased chromosomes, improves completeness by filling gaps, corrects scaffolding, and correctly phases highly heterozygous, complex regions.

## Introduction

Affordable high-throughput DNA sequencing and novel assembly tools have made high-quality genome assemblies and genome research attainable and abundant. Long-read DNA sequencing technologies, like those developed by Oxford Nanopore Technologies and Pacific Biosciences, are now the preferred methods for reference genome sequencing. The assemblies produced using these technologies are more contiguous and complete than assemblies constructed using short sequencing reads and better represent repetitive content (1–4). Another important advantage of long-read sequencing is the ability to generate phased diploid assemblies. Previously, genome complexity due to heterozygosity was typically handled by generating a haploid representation of a diploid genome either by collapsing heterozygous sites into a consensus sequence or by including only one allele’s sequence (5–10).

Partially phased assemblies have revealed genomic complexities that were inaccessible in previous haploid representations, such as haplotype-specific structural variation events, trait-associated alleles, and allele-specific gene expression and methylation (11–16). However, phasing of diploid assemblies remains challenging for complex genomes. High heterozygosity and repetitive content often prevent phasing in diploid regions. This inflates the primary assembly (17, 18) and can impair scaffolding procedures that use the primary assembly as input.

Hybrid approaches that integrate additional independent data, such as optical maps or chromatin structure, help scaffold draft genome assemblies up to full-length chromosomes (19–22). Several genetic map-based and reference-guided scaffolding tools have been developed (23–26). However, these tools do not use the relationship between haplotypes to aid the assembly process. Consequently, constructing chromosome-scale pseudomolecules using these tools relies on the phasing accuracy of the draft genome, the density of genetic map markers, or similarity to a related species’ genome (24, 26, 27). Though quality control is an integral part of the assembly procedure, the relationship between haplotypes is never included in quality control processes.

Here, we present HaploSync, an open-source package that scaffolds, refines, and fully phases diploid and chromosome-anchored genomes. HaploSync leverages the relationship between haplotypes to improve the quality and accuracy of assemblies, separates haplotypes while reconstructing chromosome-scale pseudomolecule sequences, and recovers a location for genomic regions that cannot be placed during other assembly steps. Quality controls are implemented at each step to check for and correct assembly errors. HaploSync was benchmarked using five diploid species with different levels of heterozygosity from the plant, animal, and fungal kingdoms. For each species, HaploSync delivered a completely phased, chromosome-scaled genome with a quality comparable to the assemblies considered as references for each species. HaploSync, its manual, and tutorials for its use are freely available at https://github.com/andreaminio/haplosync.

## Materials and methods

HaploSync has six modules: HaploSplit, HaploDup, HaploBreak, HaploFill, HaploMake, and HaploMap. The overall HaploSync workflow is summarized in Fig. 1. HaploSync accepts draft genome sequences or assembled pseudomolecules as input, preferably with minimally collapsed heterozygous sequences and no haploid consensus sequences. Allele phasing is unnecessary *a priori*. The tool is applicable to conventional haploid and diploid-aware assemblies.

**Fig. 1:**
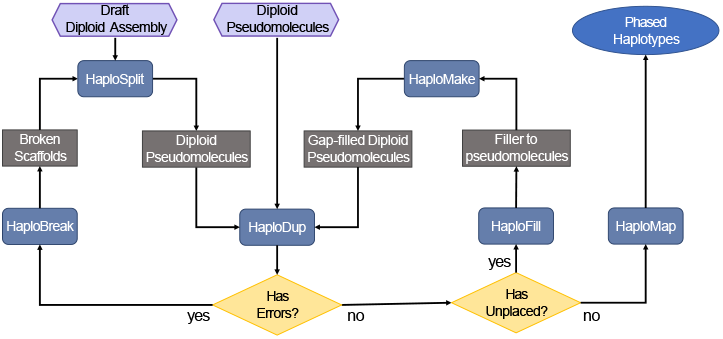
The HaploSync pipeline builds and refines haploid and diploid genome assemblies. The diploid-aware pipeline can deliver fully phased diploid pseudomolecules using a draft diploid assembly or diploid pseudomolecules as input. If draft sequences are used, Haplosplit first separates the haplotypes into two pseudomolecule sets. Pseudomolecules provided by the user or reconstructed with HaploSplit, then undergo quality control with HaploDup. If errors are found, input sequences can be edited with HaploBreak prior to rebuilding the pseudomolecules with HaploSplit. If no errors are detected and there are unplaced sequences, the pseudomolecule undergoes gap-filling with HaploFill. After each filling iteration, quality control can be performed with HaploDup. Finally, HaploMap can be used to identify colinear regions between pseudomolecules.

### A. HaploSplit

HaploSplit uses external information to associate draft assembly sequences with original chromosomes, then sorts and orients them in pseudomolecules using directed adjacency networks. Alternative sequences are detected and segregated in two different haplotypes and, if the external information relates to a chromosome, HaploSplit delivers chromosome-scale scaffolds.

External information can be a genetic map composed of sorted unique genomic markers (Fig. 2) and/or the genome assembly of a closely related species (Supplemental figure 1). When both types of information are used in hybrid mode, the genetic map is used as primary information to generate draft diploid pseudomolecules. The guide genome is used subsequently when marker information is insufficient. Phasing information between the alternative alleles is not needed *a priori*; HaploSplit will detect the existing relationship between haplotypes and phase them. The tool is capable of handling diploid assemblies lacking phasing information as well as diploid assemblies with inflated primary assemblies due to erroneous phasing. However, if the relationship between input sequences is known, it can be supplied to HaploSplit as a constraint to guide the reconstruction. For example, allelic information can be given to avoid placing primary contigs and haplotigs in the same haplotype.

**Fig. 2:**
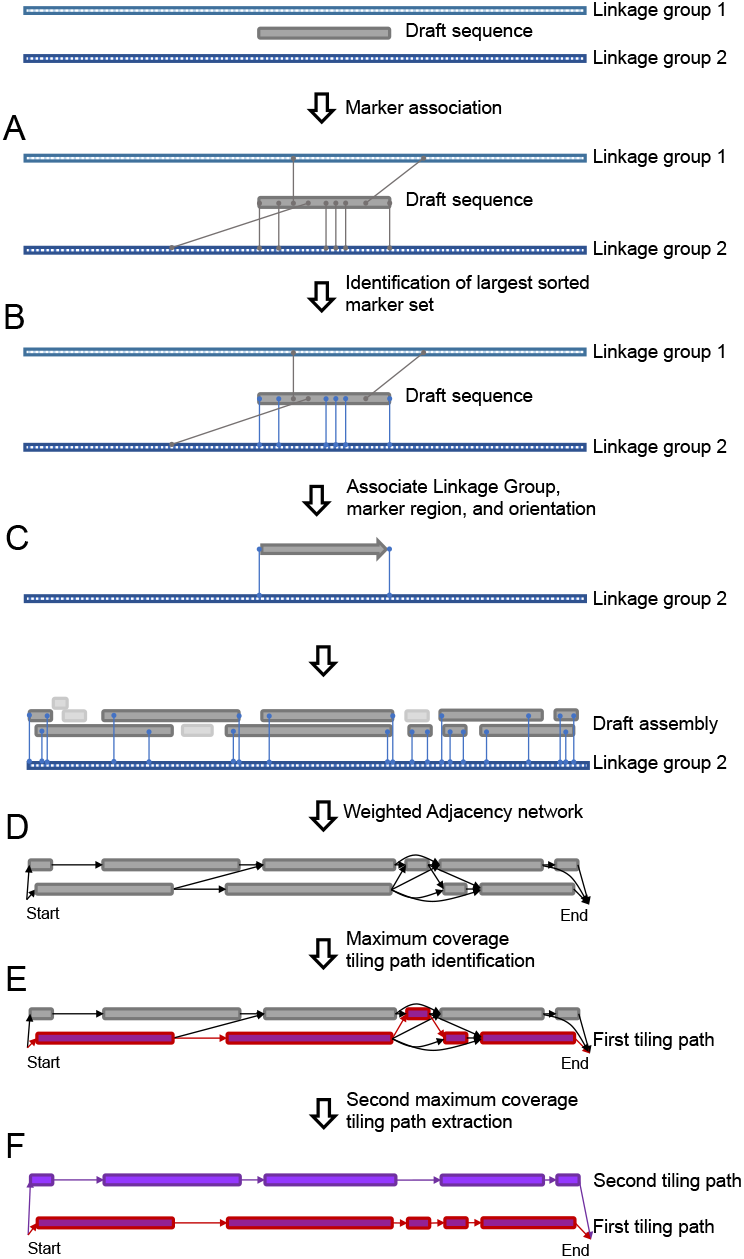
The HaploSplit procedure using genetic markers as input. A) The procedure identifies marker positions in the draft sequences. B) The longest sorted set of markers is identified for each draft sequence. C) Each sequence is assigned to a unique genomic region in the map (linkage group) and oriented. D) A directed adjacency network of non-overlapping sequences is built for each linkage group connecting all sequences with no overlapping ranges of genetic markers. Sequences sharing markers are placed in separate network paths. E) The tiling path that maximizes the number of covered markers is selected for the first haplotype. F) Sequences belonging to the first haplotype are removed from the adjacency network and the second-best tiling path is used to scaffold the second haplotype.

If a genetic map is given as external evidence, HaploSplit first assesses markers’ uniqueness and congruence in the assembly. Markers present at three or more locations in draft sequences and markers present twice in the same draft sequence, are considered unreliable and are excluded from further analysis. For each sequence containing an unreliable marker, HaploSplit produces a report containing layered interactive plots (Supplemental figure 2), including the sequence’s self-alignment, the position of reliable and duplicated genetic markers, and the copy number of annotated genes if gene annotation is available. If the input draft sequence is a scaffold, its composition in terms of legacy contigs is also included. These plots can be used to investigate the source of marker duplication within a draft sequence and to correct it using either HaploBreak (see below) or a constraint file. After identifying the genetic markers that are reliable for scaffolding, HaploSplit assigns draft sequences to a chromosome based on their largest set of consecutive markers (Fig. 2B), with their orientation based on markers’ order (Fig. 2C). If marker order does not adequately define sequence orientation (e.g., only one marker is present), the sequence is aligned and oriented based on the alternative haplotype (i.e. the sequence sharing the same marker). Once each draft sequence is assigned unambiguously to a chromosome, a directed, weighted adjacency network is created for each chromosome (Fig. 2D). Directed edges are created for each draft sequence with a weight based on the number of markers composing the sequence. Directed edges with zero weight are created to connect sequences without any common genetic marker ranges. Then, two haplotypes for each genomic region are split into different network paths. The tiling path that maximizes the number of genetic markers is used to scaffold the first haplotype and its draft sequences are removed from the adjacency network (Fig. 2E). The second-best tiling path is selected from the remaining sequences in the network and is scaffolded into the second haplotype (Fig. 2F).

If a genome is used to guide scaffolding (Supplemental figure 1), draft sequences are aligned on all guide genome sequences with Minimap2 (28). Local alignments are used to generate a directed weighted adjacency network for the query draft sequence and each guide genome sequence. Each draft sequence is associated with the guide sequence with which it shares the highest identity. Directed edges are created for each draft sequence with a significant alignment on the guide sequence. Directed edges between non-overlapping hits are added to the network and connected with a weight of zero. For each adjacency network, the tiling path maximizing the number of matching bases between the draft sequences and the guide sequence is used to build the first haplotype. The second haplotype is then scaffolded using the second best path.

When a genetic map and a guide genome are used in hybrid mode, the genetic map is used as primary information to generate draft diploid (Supplemental figure 1). The draft pseudomolecules and unplaced draft sequences are aligned to the guide genome. Then, an adjacency network is created for each guide sequence using the draft sequences composing each draft pseudomolecule and the unplaced draft sequences that do not significantly overlap the alignment of the draft pseudomolecules. The two tiling paths with the highest identity with the guide sequence are used for scaffolding the two haplotypes.

HaploSplit permits diverse, user-defined relationships between sequences to constrain and/or fine-tune scaffolding. For example, the relationship between the haplotigs and primary sequence defined by a sequence assembler like Falcon Unzip can be used to maintain consistency across alternative sequences. Similarly, a list of sequences in specific linkage groups can be given to guide their placement in pseudomolecule scaffolds.

### B. HaploDup

HaploDup (Fig. 3 and Supplemental figure 3) exerts diploid-aware quality control over pseudomolecule sequences. HaploDup generates multiple sets of interactive plots that allow the user to identify misassemblies and expose conflicts that prevent correct sequence placement. Misassemblies can be caused by erroneous hybrid scaffolding (Supplemental figure 2 Supplemental figure 3 Supplemental figure 4), a lack of colinearity information with the guide genome (Supplemental figure 5), or an incorrect sorting of genetic markers (Supplemental figure 6). Misassemblies can be inherited by downstream assembly steps if not corrected (Fig. 3 A).

**Fig. 3:**
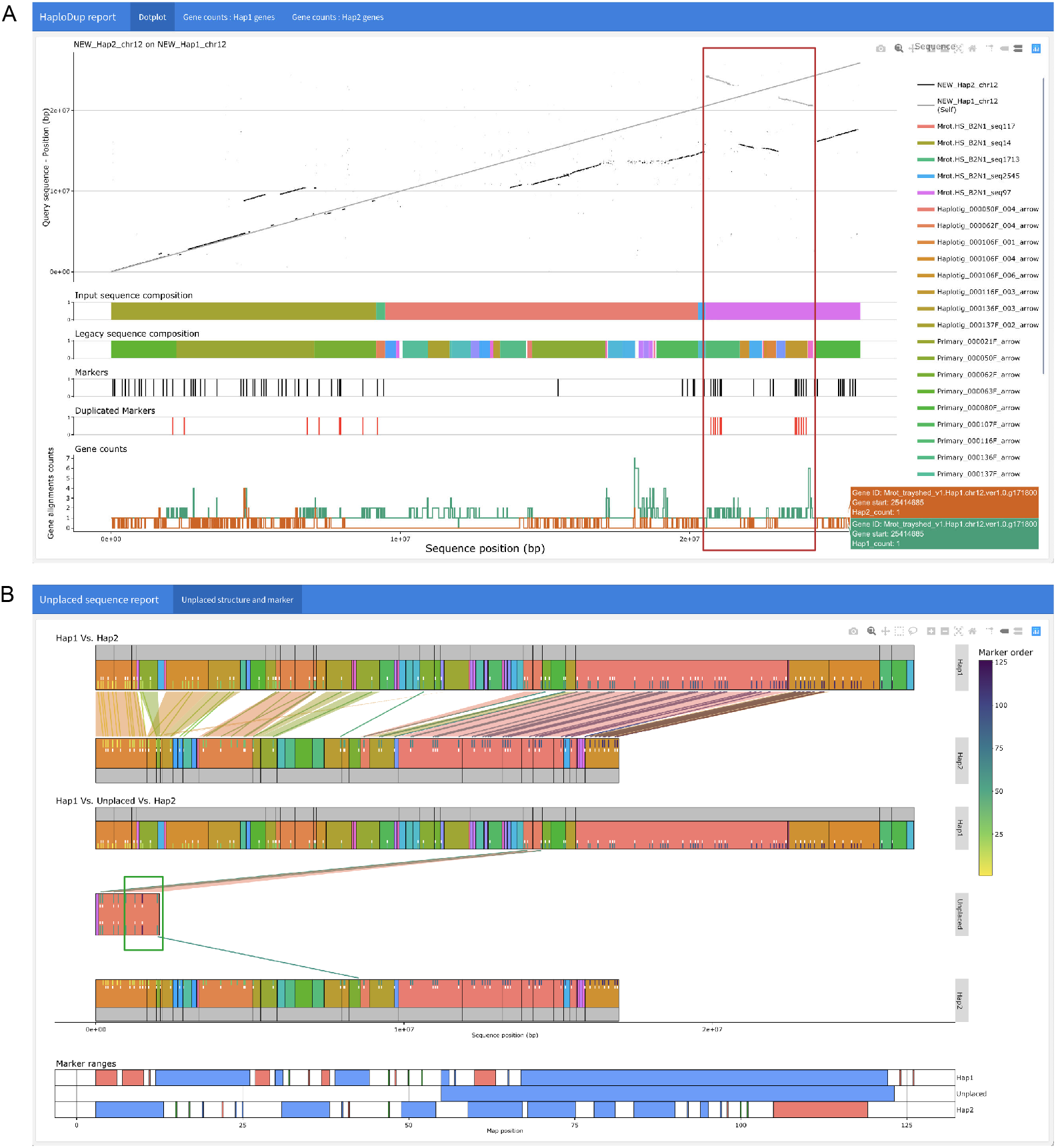
Example of HaploDup’s interactive reports. The figure reports two static screenshots exemplifying HaploDup interactive output. A) Assembly quality control of *M. rotundifolia* chromosome 12 Haplotype 1: whole-sequence alignment of both alternative haplotypes on Haplotype 1, legacy contig and hybrid scaffold composition of Haplotype 1, position of the genetic markers and the duplicated markers in Haplotype 1, number of significant alignment(s) per gene of Haplotype 1 in each alternative haplotype. In this example, the composition in legacy contigs and position of duplicated markers indicate that both alleles (primary contig and haplotig) and both marker copies were placed in a hybrid scaffold (red box). B) Unplaced sequence quality control: Marker content is compared between pseudomolecules and unplaced sequences to evaluate conditions that prevent the inclusion of a specific unplaced sequence. Color-coding is used for better contextualization. Markers are color-coded based on their order in the map. The structure of pseudomolecules and unplaced sequences are represented with color-coded blocks. Blocks identify the composition in terms of draft assembly sequences, color coding is used to show the existing relationships between the composing sequences (e.g., primary to haplotig relationships). In this example, the presence of a marker (green box, the dark violet marker on the left of the contig) in the unplaced sequences far from its expected position on the map extends the expected coverage of the map to the end of the linkage group and prevents placement in any haplotype scaffold.

To identify misassemblies and help plan a correction strategy, HaploDup compares pseudomolecule sequences, integrates structural (e.g., contigs and scaffolds) and feature (e.g., markers and genes) information, and produces interactive plots (Fig. 3). Two kinds of plots are generated. The first compares alternative haplotypes (Fig. 3 A). The second visualizes unplaced sequences with sufficient information to be placed but are currently unplaced among scaffolds because of incompatibility with other sequences; these are compared to the two alternative pseudomolecules (Fig. 3 B).

#### B.1. Alternative haplotype comparison

HaploDup produces a report for each alternative haplotype of each linkage group (Fig. 3A). The report includes layered plots: i) alignment of the two alternative haplotypes on the target haplotype; ii) the target sequence structure, with two lines of sequences at most (if available); iii) marker position and duplication status (if available).

The dotplot is essential for visualizing colinear regions within and between pseudomolecules. Duplications, deletions, and translocations can be spotted by overlaying both haplotypes’ alignments. If this information is intersected with the structure of input contigs or scaffolds, then it is possible to determine whether these peculiarities are real or are technical errors. For example, a region duplicated in one haplotype and deleted in the other may indicate that both alleles were placed in the same scaffold instead of one placed in each haplotype (red box in Fig. 3). Genetic markers and genes’ positions also help identify assembly errors. Genetic markers that are duplicated within the genome assembly are indicative of misplaced alleles. When a gene annotation is available, HaploDup counts significant alignments (>80% coverage and identity) of each CDS on its pseudomolecule of origin and on the alternative haplotype. This is useful for spotting fused haplotypes when the whole genome dotplot lacks resolution. An unbalanced number of gene copies between haplotypes in a given region can indicate a deficit of information or a duplication error. With these plots, the user can identify misassembled regions. Misassemblies can be solved by providing either a list of the breakpoint coordinates of the misplaced sequences to HaploBreak or a constraint file to HaploSplit.

#### B.2. Comparison of unplaced sequences with the two haplotypes of each pseudomolecule

HaploDup uses external information to compare unplaced sequences to related pseudomolecules (Fig. 3B). The plot reports: i) a comparison of associated pseudomolecules structures in terms of markers and sequence content. Structure is reported on two levels (scaffolds input to HaploSplit and their composition in terms of legacy contigs) when the requisite information is available; ii) a comparison of the unplaced sequence to the associated pseudomolecules in terms of markers and sequence content at two levels (scaffolds input to HaploSplit and their composition in terms of legacy contigs); iii) a comparison of the ranges of markers covered by the unplaced sequence and the ranges covered by the draft sequences composing the pseudomolecules.

Markers and their relationship to sequences can be visualized. Markers can be color-coded based on order. This plot helps resolve conflicts that prevent sequence placement into linkage groups. In Fig. 3B and Supplemental figure 6, for example, a distal marker is incorrectly ordered inside an unplaced sequence. This triggered its exclusion from any of the pseudomolecules. Once fixed, the sequence will be placed.

### C. HaploBreak

HaploBreak (Supplemental figure 7) automatically searches for and breaks sequences at the nearest known junction or at the nearest gap. The coordinates of breakpoint pairs are given by the user to estimate where sequences should be broken to correct scaffolding errors. If a scaffolding structure is supplied by the user, these junctions are prioritized to be broken. If a pair of breakpoints leads to two distinct scaffolding junctions, the original sequences reported between the two junctions will be excluded from the tiling path. If either breakpoint in a pair is associated with a sequence instead of a junction, the corresponding original sequence is broken on the nearest gap (i.e. stretch of “N” characters between two contigs). For each pair of breakpoint coordinates queried by the user, HaploBreak will do the following procedure: (i) search for scaffolding junctions closest to the two coordinates. If a junction is found within the defined search limits, it is associated with the breakpoint, else the original sequences are searched for the closest gap (i.e. a region of “N” characters), (ii) break the sequence. If the pair of coordinates is associated with two distinct scaffolding junctions (or one junction and the end of an input sequence), the original sequence between them is classified as misplaced (i.e. “unwanted” in that tiling path). If one or both breakpoints is associated with a gap in the original sequence, the sequence is broken at the gap position.

### D. HaploFill

A reference-independent approach, HaploFill (Supplemental figure 8) uses the relationship between homologous pseudomolecule scaffolds to improve the assembly’s completeness by integrating unplaced sequences where scaffolding gaps occur. Gaps are created during scaffolding procedures when adjacent regions in the pseudomolecule are assembled in separate sequences and lack sufficient information to connect them. Instead, a gap (i.e. stretches of “N” characters) is inserted as placeholder. When multiple scaffolding procedures are performed, gaps defined in previous iterations are inherited in the subsequent steps. HaploFill uses several reference-independent strategies to identify the specific kind of gap and the correct filler sequences.

A gap in a scaffold may occur when there is insufficient reliable information to identify the correct sequence for the region. This can happen when there are a lack of digestion sites in optical maps, a shortage of markers for HaploSplit, or when multiple alternative sequences are linked with proximity ligation data (e.g., mate-pair library, HiC libraries). A gap may also occur in a scaffold when the sequence is unavailable for placement. This can occur if it was not assembled or if one consensus sequence was produced from multiple genomic loci (e.g., repeats). This might also happen in diploid assemblies at homozygous regions where no alternative sequence is produced.

HaploFill is designed to recover gap information by comparing the gap region to the sequence present in the alternative haplotype. First, unplaced sequences are searched for the missing constituent. If no suitable candidate is found, the gap is filled using the alternative allele’s sequence.

HaploFill does the following steps. First, HaploFill will try to determine the ploidy of each region using sequencing coverage information: (i) align long or short sequencing reads onto each haplotype separately and calculate the base coverage along each pseudomolecule using Bedtools (29); (ii) calculate the expected haploid depth of coverage with a Savitzky–Golay filter for each pseudomolecule, excluding annotated repetitive regions; (iii) classify each region of the genome as uncovered, haploid, diploid, and repetitive based on the ratio between the depth of coverage and the expected haploid depth of coverage. Thresholds can be defined by the user. For each gap, HaploFill extracts the region upstream and downstream of the gap and the corresponding regions on the alternative haplotype to build support sequences that will assist the search for filler.

If the alternative region is reliably diploid (i.e. neither repetitive nor extensively gapped on the opposite haplotype) HaploFill will (i) create a hybrid support sequence made of the regions flanking the gap and the regions corresponding to the gap on the alternative haplotype, (ii) create an alternative support region made of the regions that correspond to and flank the gap on the alternative haplotype. If the region that corresponds to the gap on the alternative haplotype is highly repetitive or gapped, HaploFill will create two alternative support sequences made of the regions flanking the gap on the two haplotypes.

HaploFill will then search for gap filler among the unplaced sequences. To do this, HaploFill will first map unplaced sequences onto all the support regions with Nucmer (30). Unplaced sequences are assigned globally in a 1-to-1 relationship to supporting sequences. Pairings are ranked based on the bases that match non-repetitive portions of the support sequence and the whole support sequence. Then, the best filler is assigned to the gap. Filler priority is given to the hybrid support region filler, followed by the alternative support region, and then to the gapped support regions. If no filler can be validated to cover the gap but the corresponding region is classified as diploid based on sequencing coverage, the region is assumed to be homozygous. In this scenario, the region on the alternative haplotype corresponding to the gap is used as a filler. Like HaploSplit, HaploFill allows a wide range of user defined relationships between sequences to fine tune the filler selection procedure. For example, the relationship between the primary and haplotigs can be used to consistently place alternative sequences.

### E. HaploMake

HaploMake automates the conversion of sequences and annotations between different assembly versions. As input, it accepts the FASTA of the genome and a structural file (e.g., AGP files, BED, and HaploFill output files) that describes the new sequence configuration. If a gene annotation, markers, or contig structures are given, HaploMake will automatically translate their coordinates relative to the new sequence. The ends of adjacent regions in the structure files can be checked for overlaps with Nucmer (30). The coordinates of adjacent regions can be corrected by adjusting junction positions. This avoids duplicating genomic content in the final sequence and can be done without altering the gene annotation (Supplemental figure 9).

### F. HaploMap

HaploMap (Supplemental figure 10) performs a pairwise comparison between haplotypes and delivers a pairwise tiling map of colinear, non-overlapping, and non-repetitive regions between different haplotypes. Like HaploSplit, local alignments between each pair of sequences are performed with Minimap2 (28) or Nucmer (30). Hits are used to create a weighted adjacency graph for identifying a bidirectional tiling path that maximizes the identity between the two sequences. The coordinates of the colinear regions that form the bidirectional tiling path are listed in a pairwise, phased map of matching sequences.

### G. Testing datasets

HaploSync performance was tested using a wide range of species and assembly protocols (Table 1). The diploid *Candida albicans* draft assembly (31), built using PacBio reads and FalconUnzip (17), was anchored to chromosomes using the genetic map generated by (32). A diploid genome assembly of *Arabidopsis thaliana* Columbia-0 (Col-0) X Cape Verde Islands (Cvi-0) (17) was anchored using a genetic map from (33). The *Bos taurus* Angus x Brahma genome from (34) was assembled using FalconUnzip, anchored to chromosomes using the genetic map from (12), and integrated with sex chromosome information from the Integrated Bovine Map from Btau_4.0 release available from https://www.hgsc.bcm.edu/other-mammals/bovine-genome-project. To support the assembly and quality control of pseudomolecule reconstruction, the locations of unique genes from the respective reference annotations (*C. albicans* SC5314_A22, *A. thaliana* TAIR10, *B. Taurus* Btau_ARS-UCD1.2) were identified by mapping CDS sequences on primary and haplotig sequences using GMAP (ver. 2019.09.12 (35)). Unique gene models were defined by mapping CDS sequences from the reference genomes annotations on the respective reference genome sequences using GMAP (ver. 2019.09.12 (35)). All CDS mapping on multiple locations in the haploid genome were removed from the dataset. HaploFill was applied once to each of these three genomes. The *Vitis vinifera* ssp. *vinifera* cv. Cabernet Franc FPS clone 04 genome was assembled and scaffolded with PacBio reads and Dovetail HiC data (36). *Muscadinia rotundifolia* cv. Trayshed contigs were assembled with FalconUnzip in hybrid scaffolds that used BioNano NGM maps (37). A *Vitis* consensus genetic map (38) was used to anchor both genomes to chromosomes in HaploSplit and followed by several iterations of HaploFill.

**Table 1:**
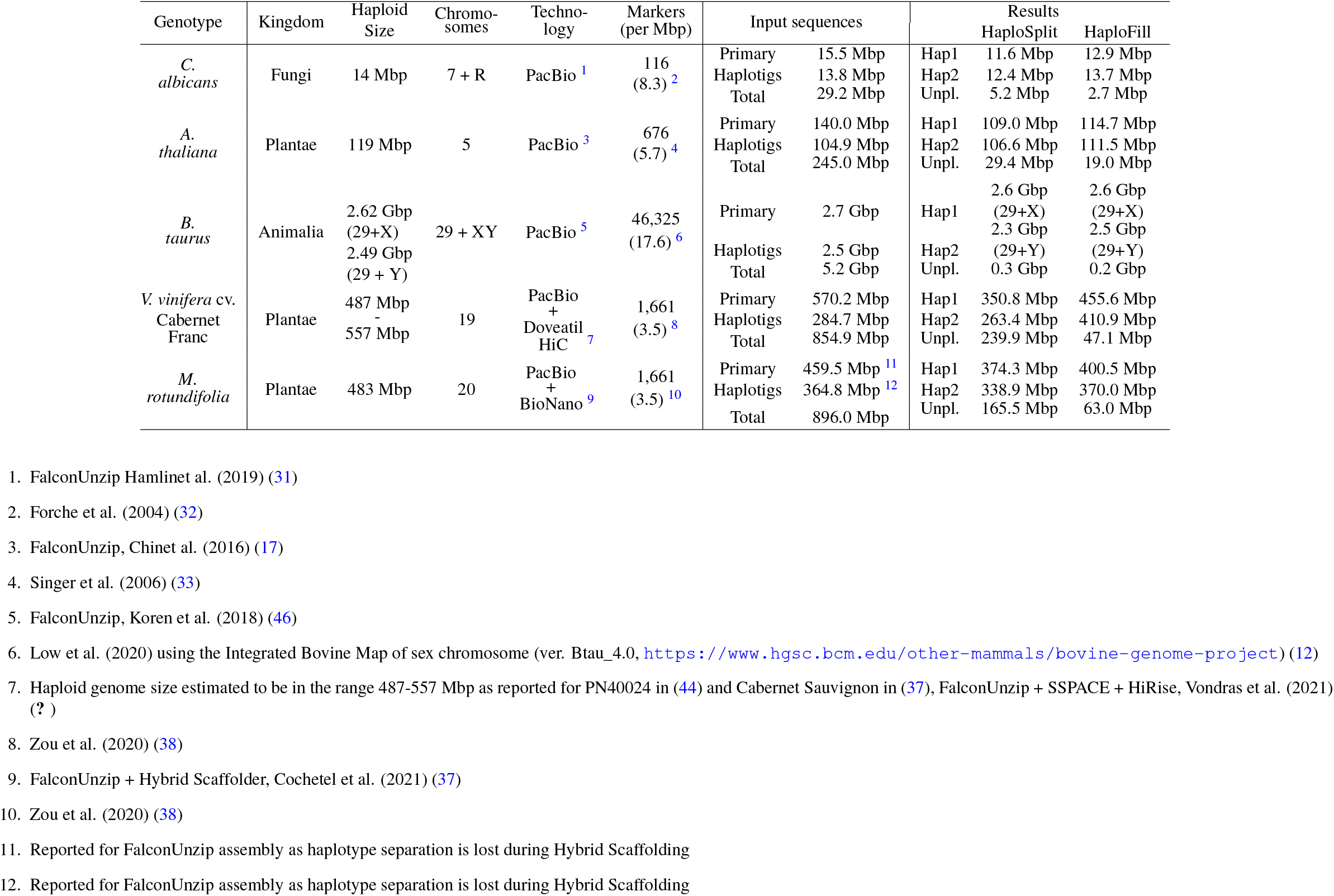
Assembly statistics

## Results and discussion

To evaluate HaploSync’s performance, five diploid species from three different kingdoms were selected. This included a *Muscadinia rotundifolia*, *Vitis vinifera*, and an F1 progeny of *Arabidopsis thaliana* (Col-0 x Cvi-0) (17), the bull *B. taurus* Angus x Brahma (12, 34), and pathogenic yeast *Candida albicans*. These species are diverse and vary in genome size, chromosome number, repeat content, and amount of heterozygosity. Long sequencing reads, genetic maps, and public reference genomes are available for those species.

### H. HaploSync adaptability to different species

HaploSync produced high-quality genomes for all five species (Table 1). The resulting assemblies were nearly twice the size of their original haploid assemblies, with 1.87X to 2.03X their gene space represented (Supplemental table 1). This indicates that most of both haplotypes were assembled separately. High-density genetic maps and highly contiguous draft assemblies enabled HaploSplit to produce high-quality pseudomolecules that differed 5.8 - 17.8% from their expected chromosome sizes. In one iteration, HaploFill increased assembly completeness and reduced the difference in length between haplotypes.

For *C. albicans*, the limited number of markers were used to anchor 2.4 Mbp of sequences to pseudomolecules in HaploSplit. The assembly had the highest share of unplaced sequences (5.2Mbp), but HaploFill recovered 17.9% of the missing genomic content in one iteration. The final pseumolecules were up to 97.9% complete. Only 231 (3.8%) of 6,079 single copy genes in the reference annotation mapping on the assembled sequences were not represented in the pseudomolecules produced by HaploSplit. This number was reduced to 182 (3.0%) in a single iteration of HaploFill. BUSCO analysis confirmed the nearly complete separation of alternative alleles with only five complete gene models found in multiple copies in Haplotype 1 (3 genes) and Haplotype 2 (2 genes) (Supplemental table 1).

With 18.6 ± 0.6 markers/Mbp, the genetic map of *B. taurus* autosomal chromosomes was the most dense out of the species used in this study. HaploSplit produced pseudomolecules almost identical in size to the ARS-UCD1.2 genome assembly (39), with Haplotype 1 pseudomolecules deviating by 0.7 ± 0.7% and Haplotype 2 by 6.7 ± 3.0% (Supplemental table 2, Supplemental figure 11). HaploFill inserted 151 Mbp, mostly in Haplotype 2 pseudomolecules, reducing missing information in Haplotype 2 pseudomolecules to 1.4 ± 1.9% of ARS-UCD1.2 chromosome sizes. For sex chromosomes, only a genetic map of the X chromosome with low marker density was available (2.1 markers/Mbp, assembly ver. Btau_4.0 available at https://www.hgsc.bcm.edu/other-mammals/bovine-genome-project). As a consequence, HaploSplit’s performance dropped. HaploSplit retrieved 79.8% of the expected 139 Mbp X chromosome. However, HaploFill reduced missing information to 9.7% (Supplemental table 2, Supplemental figure 11). Without markers available, the length of the Y chromosome was only 11% of its expected size (4.5 Mbp). The gene space was more complete in terms of single copy reference genes. Only 7 of 57,974 single copy CDSs mapping on the assembled sequences were not placed in the initial pseudomolecules produced with HaploSplit. This was reduced to 5 by HaploFill. BUSCO analysis confirmed the completeness and the separation of the alleles, with 92.5% complete gene models found in Haplotype 1 (1.3% in multiple copies) and 86.8% in Haplotype 2 (1.2% multiple copies) (Supplemental table 1).

In plants, the high level of polymorphism and structural variation between haplotypes make assembly and phasing challenging (17).

The high level of heterozygosity in the *A. thaliana* accession used to test HaploSync is caused by sequence variation between its parents, Col-0 and Cvi-0. This led to a primary assembly 17% longer and haplotigs 11.8% shorter (17) than the haploid reference genome. After Haplosync, the two sets of pseudomolecules differed by 3.6% and 6.3% from the haploid reference genome size. This supports the tool’s ability to phase duplicated primary content between haplotypes. When gene space completeness was estimated using single-copy genes in the reference annotation, similar results were obtained. The amount of single copy CDSs mapping on the assembled sequences represent the 99.7% of the entire dataset (34,741 out of 34,854). After HaploSplit, unplaced sequences included 1,966 putative loci (5.7%). Of these, 261 (1.1%) were missing from the pseudomolecules. HaploFill further increased the completeness of the pseudomolecules to include 98.1% and 95.8% of the gene space in the two haplotypes. This reduced the putative, single copy CDS loci among unplaced sequences to only 123. Over 97% complete BUSCO gene models were complete in Haplotype 1 and the Haplotype 2, with only 1.3% and 1.5% in multiple copies, respectively (Supplemental table 1).

*Vitis* species can be 12% heterozygous (40). Assemblies of the species can exhibit extensive loss of phase between primary sequences and associated haplotigs (17, 18, 41–43). In Cabernet Franc, for example, the primary assembly is inflated by 18.8% and haplotigs are 40.7% shorter than the expected haploid genome size. HaploSync was able to overcome these limitations for both species and placed over 93.0% of the sequences in phased pseudomolecules that were no more than 9.8% different in size. HaploSplit also automatically placed and correctly phased the grape sex determining region (13) in *Muscadinia* and *Vitis* species. Using the unique CDS sequences from PN40024 as a reference for *Vitis* gene space, 1,233 (6.2%) of genes could not be placed in Cabernet Franc pseudomolecules with HaploSplit and 223 (1.4%) of genes could not be placed in *M. rotundifolia* pseudomolecules. This fraction of gene coding sequences could not be placed because of high fragmentation and low, uneven marker density that negatively affected pseudomolecule reconstruction performance. Several iterations of HaploFill reduced the number of unplaced CDSs to 0.3% for both genomes. This included 91 and 46 unique genes among unplaced sequences for Cabernet Franc and Trayshed, respectively. This highlights HaploFill’s ability to recover gene space information. Completeness and phasing of both Haplotypes was confirmed with BUSCO: 93% complete models in Haplotype 1 and 83% in Haplotype 2.

### I. HaploSync performance adaptability to different assembly procedures

HaploSync was applied to two grapes, *M. rotundifolia* cv. Trayshed (37) and *V. vinifera* cv. Cabernet Franc (36), to assess its adaptability to genomes assemblies produced using different strategies. Although contigs were produced with PacBio data and FalconUnzip for both draft assemblies, Trayshed and Cabernet Franc were scaffolded with different technologies. *M. rotundifolia* underwent hybrid scaffolding with PacBio and a NGM map, which matches optical fingerprints of DNA molecules with assembled sequences digested *in silico* with the same enzyme. Gaps were introduced where there was a low density of digestion sites. Systematic errors were observed at highly repetitive and heterozygous regions, including the RUN1/RPV1 locus on chromosome 12 (Supplemental figure 2). The differential expansion of TIR-NBS-LRR genes between haplotypes (37) may have caused their fusion in the same scaffold. These issues affected 50 hybrid scaffolds (326.2 Mbp), required correction, and were easily found with HaploDup. For Cabernet Franc, scaffolding was performed using HiC data that produced chimeric scaffolds due to the presence of diploid information in the primary assembly. Both haplotypes of 108 of scaffolds (449 Mbp) were included in the same assembled sequence (Supplemental figure 4).

After scaffold correction, both genome assemblies were anchored to chromosomes using a *Vitis* consensus genetic map (38). Low specificity and marker density (3.5 markers/Mbp) affected the construction of pseudomolecules by HaploSplit and negatively affected HaploSync’s performance. Cabernet Franc was most affected, with only 350.8 Mbp and 263.4 Mbp placed on Haplotype 1 and Haplotype 2, respectively (i.e. 75% and 55% of the reference haploid genome). Unpleaceable sequences were nearly half of Cabernet Franc’s expected haploid genome size (240Mbp). Trayshed’s assembly was more complete; Haplotype 1 and Haplotype 2 assemblies were 374.3 Mbp and 338.8 Mbp long, respectively.

Three iterations of HaploFill were performed on Cabernet Franc’s assembly. Each iteration reduced unplaced sequences by nearly one half (Supplemental figure 12). The final Cabernet Franc pseudomolecules were 456 Mbp (Haplotype 1) and a 411 Mbp (Haplotype 2). Afterwards, 47 Mbp (5.4%) of sequences remained unplaced. In contrast, only two iterations of HaploFill were sufficient to leave just 8% of Trayshed sequences unplaced. Haplotype 1 and Haplotype 2 of Trayshed’s pseudomolecules were 400 Mbp and 370 Mbp, respectively. The total sizes of both haplotypes in both chromosome-scale assemblies were similar to their expected haploid reference genome sizes (44, 45) and to Cabernet Sauvignon’s haplotypes (459 Mbp and 449 Mbp, respectively) (13).

### J. HaploSync performance assessment

The performance of different HaploSync tools, in terms of result quality and processing time, are influenced by multiple factors. Unsurprisingly, the genome size and the number of linkage groups affect all assembly phases and the duration of alignment procedures. For HaploDup, HaploFill, HaploMap, HaploBreak, and HaploMake, genome size determines the size of the output and how long alignments take to complete, which can constitute over 90% of the computational time. The number of linkage groups exponentially increase the number of comparisons and plots needed. For example, HaploDup required 40 hours to process *B. taurus*, which has a 2.6 Gbp haploid genome size in 30 linkage groups and is the largest dataset used in this study. Nearly 15 of these hours were consumed for alignments between sequences while using 24 cores. *C. albicans* is the smallest dataset, with 14 Mbp in 8 linkage groups. In contrast to *B. taurus*, the same procedure required 75 minutes, with only 5 minutes dedicated to mapping.

HaploFill performance is also affected by the number of phased genomic sequences in the pseudomolecules. Alternative pseudomolecules are the backbone that enable the algorithm to retrieve gap filling information. The completeness of the pseudomolecules directly affects the amount of information usable as support for sequence placement. Unplaced sequences are information that might be recovered. The workflows adopted for *A. thaliana* and for the *Vitis* genotypes were selected based on pseudomolecule completeness. The *A. thaliana* assembly had relatively low sequence fragmentation and a high density map. The pseudomolecules created for *A. thaliana* with HaploSplit were fairly complete after a single filling procedure. HaploSplit was less effective for Cabernet Franc and Trayshed because their assemblies were more fragmented and their maps were less dense. The workflow used for the grape genomes included several iterations of HaploFill to achieve highly complete pseudomolecules (Supplemental figure 12).

HaploSplit is fast. It takes between a few seconds and one minute to build the adjacency graph, traverse it, find the two best tiling paths, and report the structure of the phased pseudomolecules. In contrast, the input quality control and the alignment between the draft sequences and the guide genome in preparation for the graph creation can be time-consuming. HaploSplit result quality is affected by several factors. The disparity and incomplete representation of both alternative alleles affects the completeness of the diploid pseudomolecules produced and necessitates filling. *A. thaliana* and *B. taurus* are F1 progeny. Their considerable structural variability is captured by the FalconUnzip assembler, which reconstructs the alleles fully and separately. In contrast, Cabernet Franc and Trayshed have several homozygous regions that were assembled in a single copy and highly heterozygous regions that increased the fragmentation of the contigs by fooling the assembler into overassembling the primary sequences. This difference is reflected in HaploSplit’s results. HaploSplit was able to separate alleles and deliver a nearly complete diploid assembly of *A. thaliana* and *B. taurus*. *Vitis* required a more extensive filling procedure to recover the missing information.

HaploSplit can use a genetic map and/or a guide-genome as information to facilitate scaffolding. We tested how HaploSplit performs given different scaffolding information using *Vitis vinifera* cv. Cabernet franc cl. 04 (36). Reference genomes of closely related accessions, PN40024 (45) and Cabernet Sauvignon (13), are available. Cabernet Franc contigs were scaffolded (i) with the genetic map of (38), (ii) using the PN40024 V2 assembly or the first haplotype of Cabernet Sauvignon as guides, or (iii) using both the genetic map and a guide genome. The reference-based approach incorporated more sequences into pseudomolecules than when only a genetic map was used. As expected, the best results were obtained using Cabernet Sauvignon as a reference, which shares one allele with Cabernet Franc. This approach, however, led to overfitting of the scaffolding results to the guide. Small structural variants in long draft sequences (Supplemental figure 13 A boxes) can find a proper representation thanks to neighbouring colinear regions. Larger structural variants that encompass multiple sequences may fail to be reported correctly together. Each draft sequence location is identified independently from the others based on colinearity with guide genome, so placement is based on the structure of guide sequences rather than their actual order (Supplemental figure 13 B boxes). Moreover, gaps or the lack of information in the guide genome may impede the recovery of novel information. Only draft sequences that partially anchor within present information can be placed (Supplemental figure 13 C boxes). As a consequence, fragmented draft assemblies and the second haplotype are prone to be artificially similar to the guide genome. The hybrid approach performs better. The reconstruction of both haplotypes is more complete than the map-based approach, with the second haplotype benefiting most from this strategy (Fig. 4 A). Though no overassembly was observed, the mapping phase duplicated some alleles. Both copies of several markers occured in the same psedomolecule scaffold when Cabernet Sauvignon (5 markers) and PN40024 (4 markers) were used as guides.

**Fig. 4:**
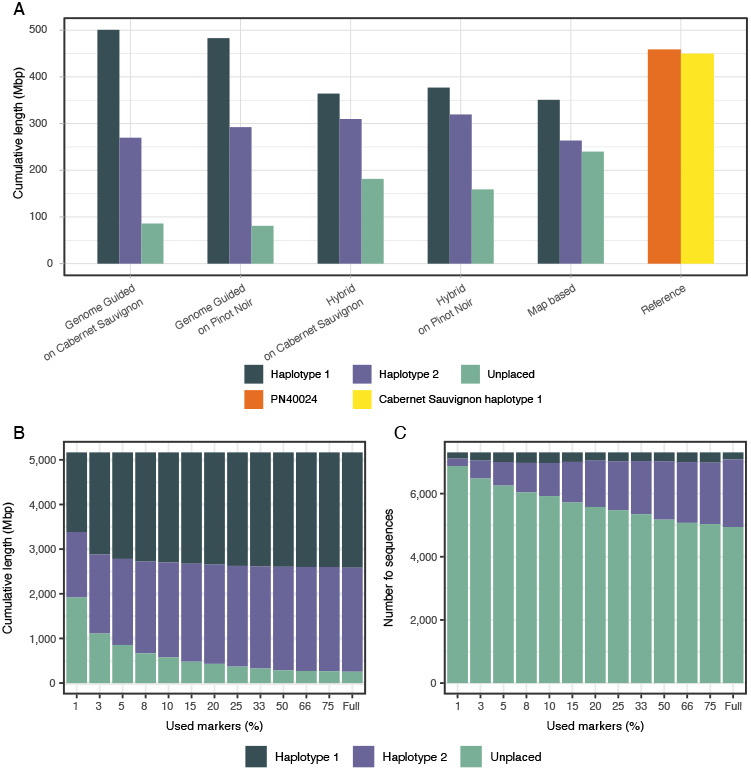
HaploSplit performance. A) The results of using different sources of external information and HaploSplit protocols for *Vitis vinifera* cv. Cabernet Franc cl. 04 (36) assembly. Map-based assembly produces the largest first haplotype, but its overassembly occurs at the expense of the second haplotype’s completeness. A map based approach is conservative and limited by the density of the markers. The hybrid approach recovers more sequences where the map is lacking information, without overassembling, and delivers a better reconstruction of both haplotypes. B) Effect of limited marker availability on overall assembly length tested on *B. taurus* Angus x Brahma (12, 34) by subsampling the genetic map. Longer sequences are more likely to contain a marker, making the first reconstructed haplotype most complete across all tests and with little variation in size. As the number of available markers increases and short sequences are included, the completeness of the second haplotype improves. C) Effect of limited marker availability on the number of placed sequences tested on *B. taurus* Angus x Brahma (12, 34) by subsampling the genetic map. Increasing the number of markers as fragmentation increases allows recruiting more sequences for scaffolding and improves completeness. Haplotype 1, with long sequences, shows little variation. In contrast, Haplotype 2 greatly benefits from increased marker density. The majority of sequences that remained unplaced are short and a small fraction of the genome’s length.

The effect of the number of reliable genetic markers on the performance of HaploSplit was tested on the genome of *B. taurus* Angus x Brahma (12, 34). The same diploid genome underwent chromosome scale reconstruction using a randomly selected subset of 479 markers (1%, 0.2 markers/Mbp) out of its available genetic map (46,323 markers, 17.6 markers/Mbp; Fig. 4 B and C, Supplemental figure 14). Unsurprisingly, the number of unplaced sequences increased to 37% of the total assembly length given lower marker density. HaploSplit found a location in pseudomolecules for 99.4% to 100% of the sequences with markers. The performance of the algorithm, in terms of completeness of the delivered pseudomolecules, is primarily influenced by input assembly fragmentation and the genetic map’s marker density. This limits the number and the sizes of the sequences with markers. The primary assembly is composed of extremely long sequences that likely contain markers and are placed even when map density is low. In contrast, Haplotigs are more fragmented and require high marker density for comparable coverage. As a result, the first haplotype assembly is more complete even with fewer markers present (Fig. 4 B and C).

In summary, the type and quality of the external guide information have a large effect on the quality of the final assembly. A guide genome aids assembly via local sequence alignments; lack of homology between sequences and repetitive regions can cause segregation errors (Supplemental figure 6), misplacements, and overfitting to the guide genome structure. Genetic maps are more conservative, with the uniqueness of markers requiring a coherent placement within a map, if at all. Moreover, errors in the map can be more easily addressed by the user than errors in the guide genome sequence. The efficiency of HaploSync relies heavily on map precision (Supplemental figure 6) and the density and evenness of its markers (Table 1).

## Conclusions

These results emphasize the importance of controlling and correcting the sequences used as input to HaploSplit to prevent scaffolding errors. Although map quality and marker density affect pseudomolecule construction by HaploSplit, HaploFill generated phased assemblies with few unplaced sequences and sizes similar to their haploid reference genomes. Sequencing technologies and assembly tools are continuously improving. HaploSync delivers assemblies with unprecedented quality and contiguity that can provide novel insight into genome structure and organization. The HaploSync suite of tools can be used to address some of the remaining impediments to genome reconstruction and improve assembly quality by taking advantage of diploid information that is readily available. HaploSync correctly and completely phases diploid genomes, reconstructs pseudomolecules by recovering missing information, and exerts quality control over the results.

## Web resources

HaploSync is freely available for download at GitHub https://github.com/andreaminio/haplosync. Instructions for installation, a full list of dependencies, a description of each tool, and tutorials are available in HaploSync’s manual (https://github.com/andreaminio/HaploSync/tree/master/manual).

## Data availability

The data used in this study are summarized in Supplemental table 3. Pseudomolecule reconstructions of *Candida albicans* NCYC4145 (31), *Arabidopsis thaliana* Columbia-0 (Col-0) X Cape Verde Islands (Cvi-0) (17), and *Bos taurus* Angus x Brahma (34) are available at Zenodo (https://zenodo.org/record/3987518, DOI 10.5281/zenodo.3987518). *Vitis vinifera* cv. Cabernet Franc cl. 04 (36) and *Muscadinia rotundifolia* cv. Trayshed (37) pseudomolecule assemblies are available at www.grapegenomics.com.

## Authors’ contributions

AM and DC conceptualized the project. AM developed the methodology and software. AM and NC performed bioinformatic analyses and tested the software. AM, NC, and AMV wrote the software manual and pipeline walk-through. AM, AMV, MM, and DC wrote the manuscript. DC secured the funding and supervised the project. All authors have read and approved the manuscript.

## Supplemental materials

**Supplemental figure 1:**
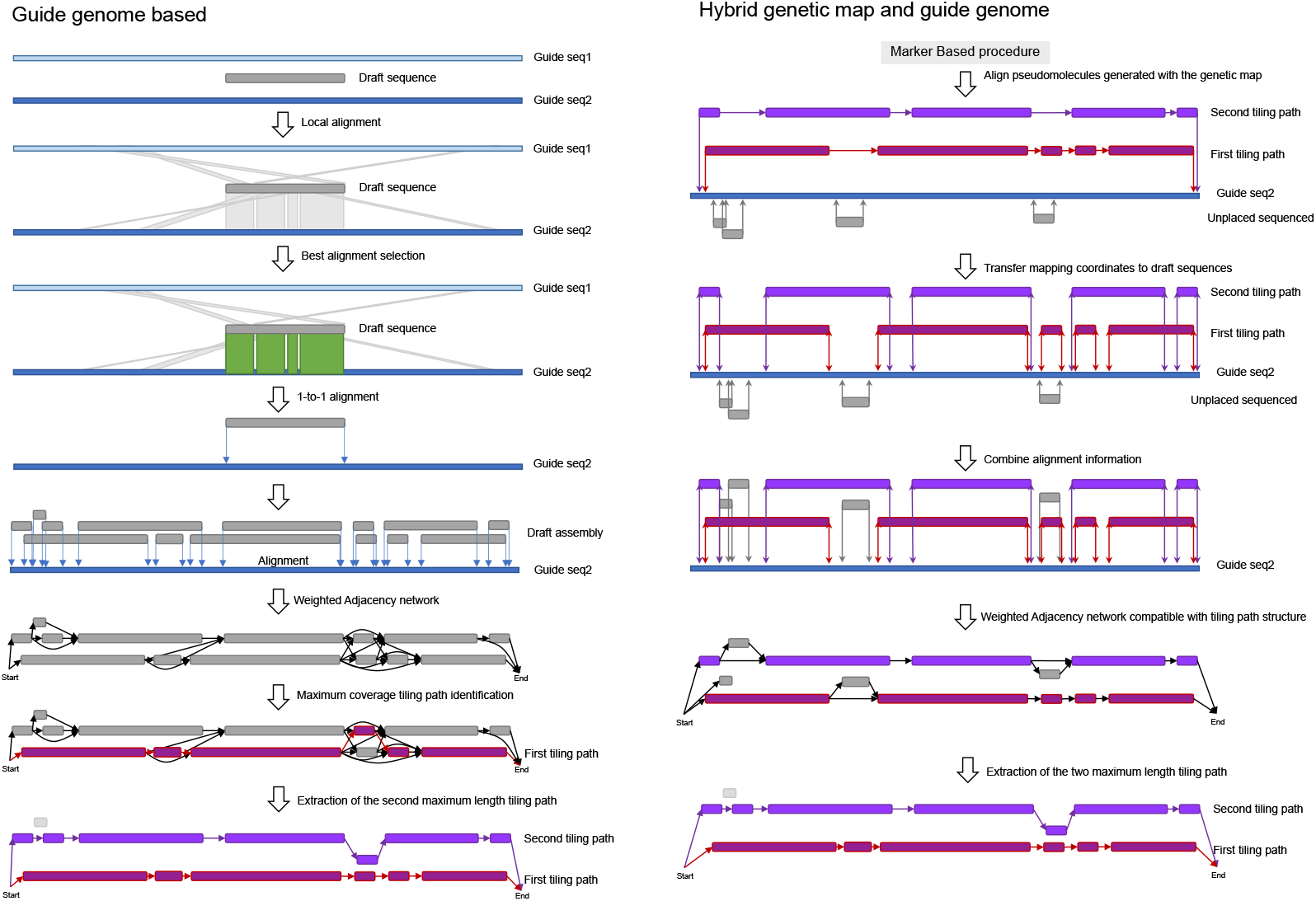
HaploSplit workflow diagrams for genome guided pseudomolecule reconstruction and hybrid reconstruction with both map and guide genome information. HaploSplit reconstructs pseudomolecules using colinearity with a guide genome to sort sequences into different haplotypes. i) Query draft sequences are aligned onto the guide genome and used to build a directed adjacency network of non-overlapping hits. ii) The tiling path that maximizes the number of identical bases between the query and guide sequence is selected. iii) A directed adjacency network of all query sequences associated with a guide sequence is created. iv) The tiling path with the highest number of identical bases with the guide sequence is used to generate the first haplotype. v) Sequences belonging to the first haplotype are removed from the adjacency network and the second-best tiling path is used to scaffold the second haplotype. In hybrid mode, after performing the reconstruction with the genomic map, i) intermediate pseudomolecules and unplaced sequences are mapped on the genome are mapped on the guide genome; ii) location of each draft input sequence composing the pseudomolecule is translated from whole sequence local alignments; iii) an adjacency graph graph of all mapped draft sequences is created; iv) the two best tiling paths are identified and final pseudomolecules are delivered.

**Supplemental figure 2:**
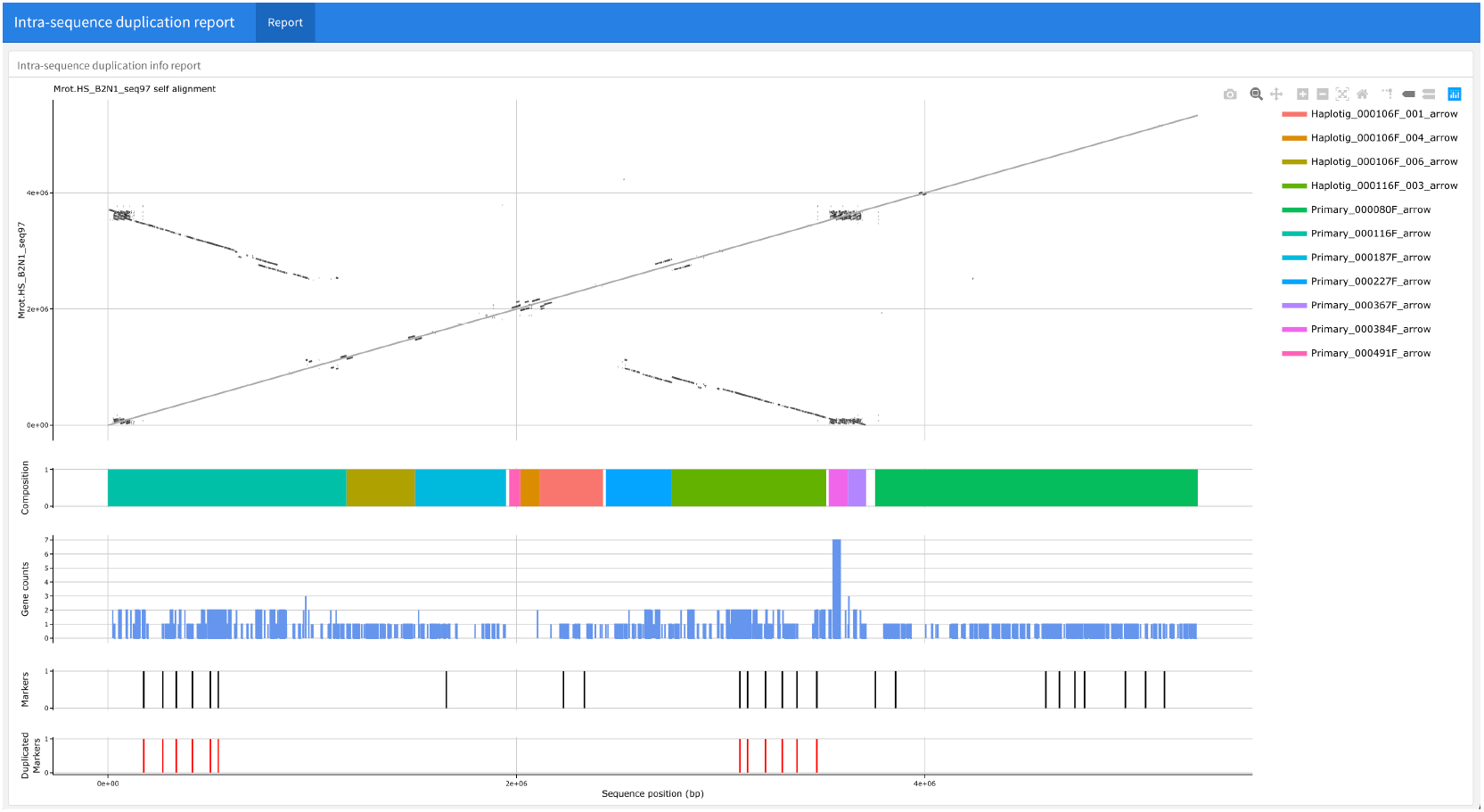
HaploSplit input QC interactive plot. Example of interactive plot generate by HaploSplit to perform a a quality control of the input sequences as a intra-sequence duplication of genetic markers has been detected. The self alignment of the sequence is performed by 1 Kbp windows to increase the resolution of duplicated regions identification. Markers present in the sequence and duplicated ones, are also reported and, when available, the sequence structure and the gene copy count tracks are layered to support the duplication analysis. The plot evince and example of systematic errors observed in correspondence to highly repetitive and heterozygous regions when FalconUnzip produced contigs undergo Hybrid Scaffolding with BioNano NGM maps. In the example, RUN1/RPV1 locus on chromosome 12 (same as Figure 3). Different expansion of TIR-NBS-LRR genes between haplotypes is the probable cause the fusion of both haplotypes of the region in the same scaffold. These issues affected 50 hybrid scaffolds (326.2Mbp) requiring correction.

**Supplemental figure 3:**
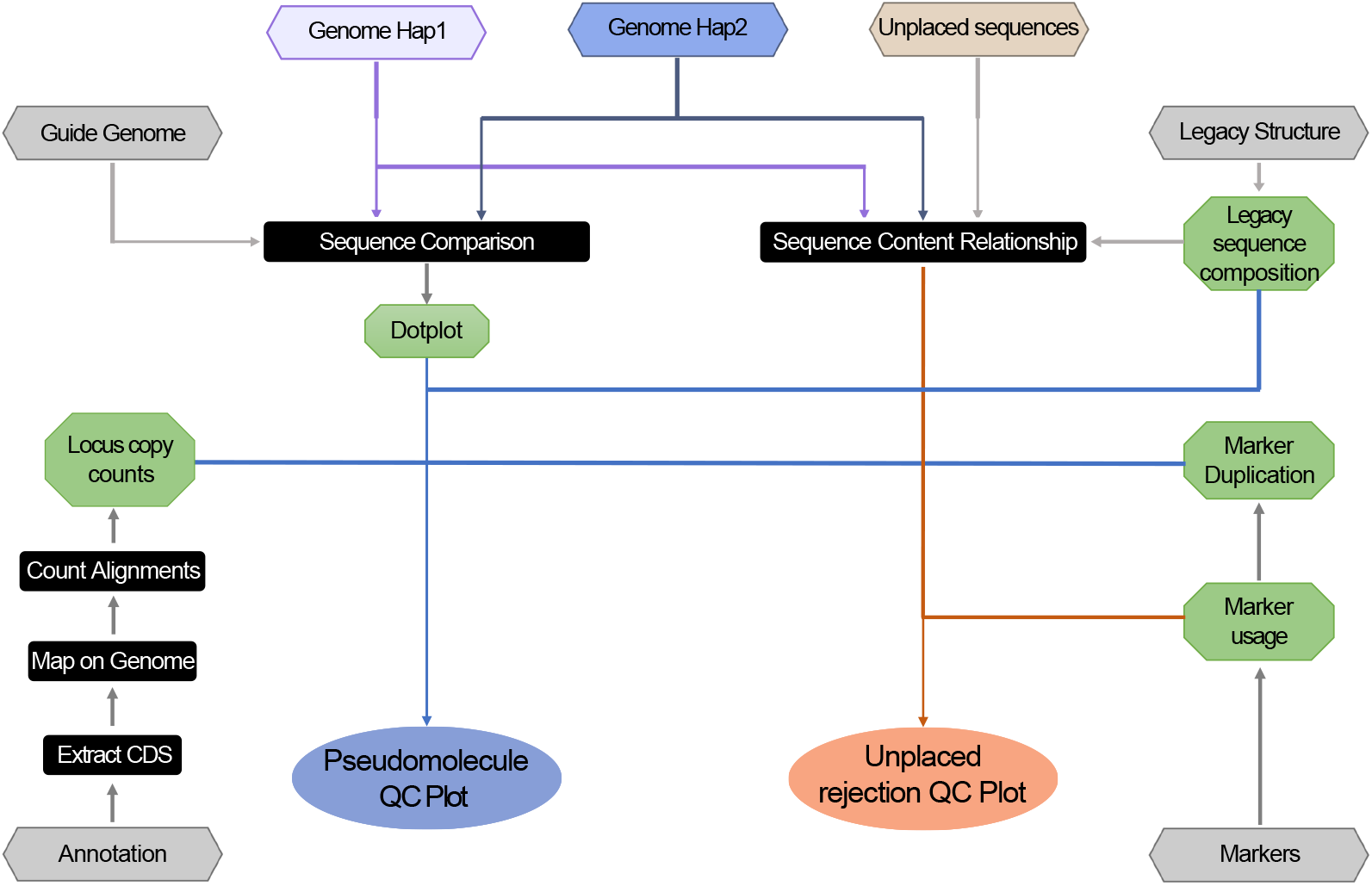
Schematic representation of HaploDup procedure. Diagram of HaploDup data usage to create Quality Control interactive plots

**Supplemental figure 4:**
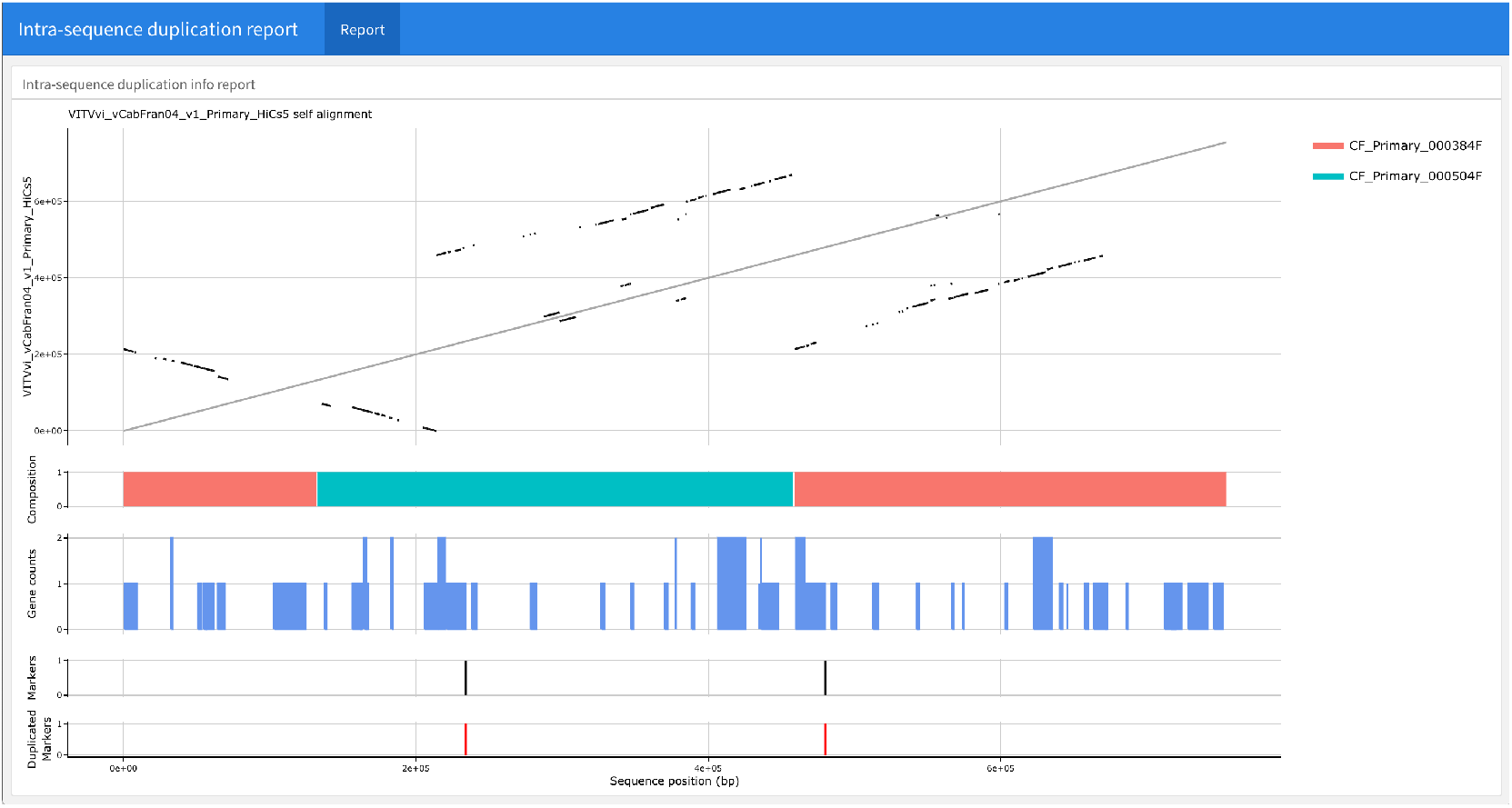
Example of assembly error due to scaffolding of primary sequences with Dovetail HiC data. Scaffolding FalconUnzip produced primary contigs with HiC data produced chimeric scaffolds. This is due to the presence of both haplotype information in the overassembled primary primary set of contigs. As a result, one haplotype is split to accommodate the alternative one inside. For Cabernet Franc, correction was required by 108 of scaffolds encompassing 449Mbp.

**Supplemental figure 5:**
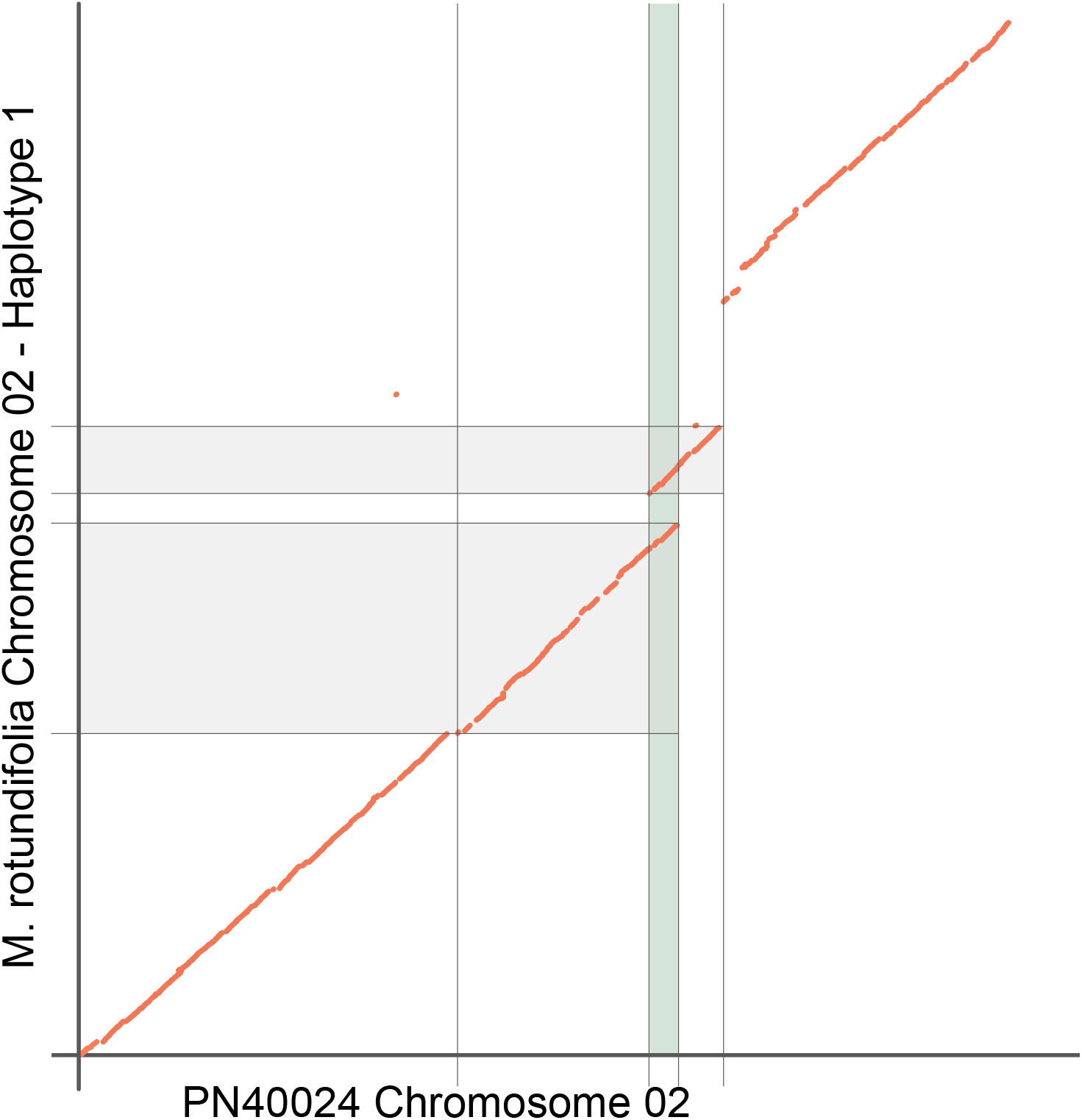
Example of errors due to lack of overlap. Lack of colinearity information in the sequences extremities in HaploSplit may not allow a proper identification of homologous sequences. The example here reported shows an example. Plot was made using Nucmer and reporting all alignment hits without filtering, It is evident that the inner extremity of the two inserted sequences (in grey) do actually represent the same genomic region (in green). Due to the alignment with Minimap on the genome, however, the projection of the sequences on the guide genome did not report the conflicting overlap, thus allowing the placement of both sequences in the same pseudomolecule scaffold.

**Supplemental figure 6:**
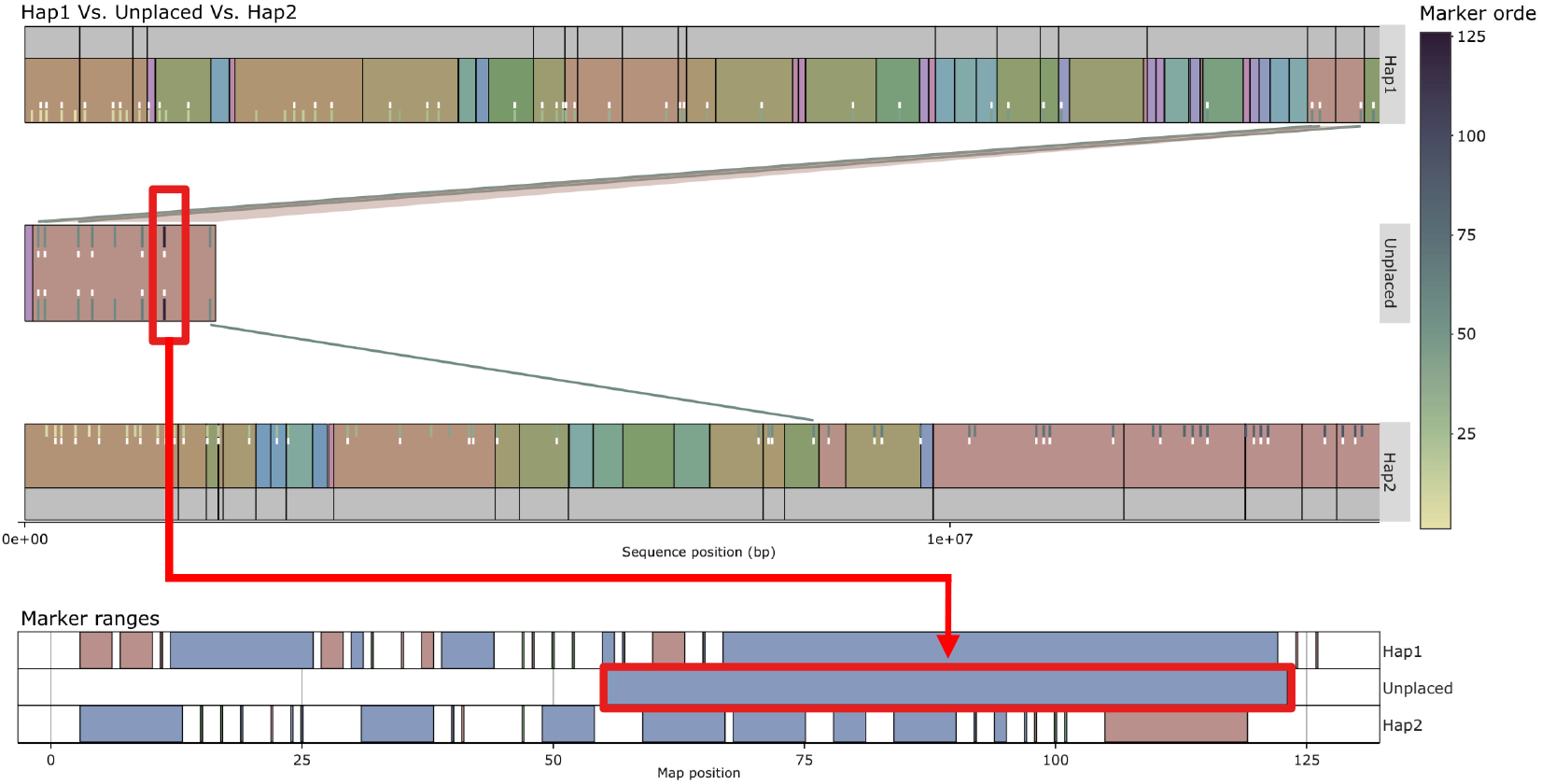
Example of errors due to marker sorting errors. Erroneous sorting order of markers may not allow placement of a sequence because of conflicting map coverage. The presence of the marker at position #123 of the map in the contig caused HaploSync to overestimate the extension of the range covered markers, making it impossible to place the draft sequence in any haplotype.

**Supplemental figure 7:**
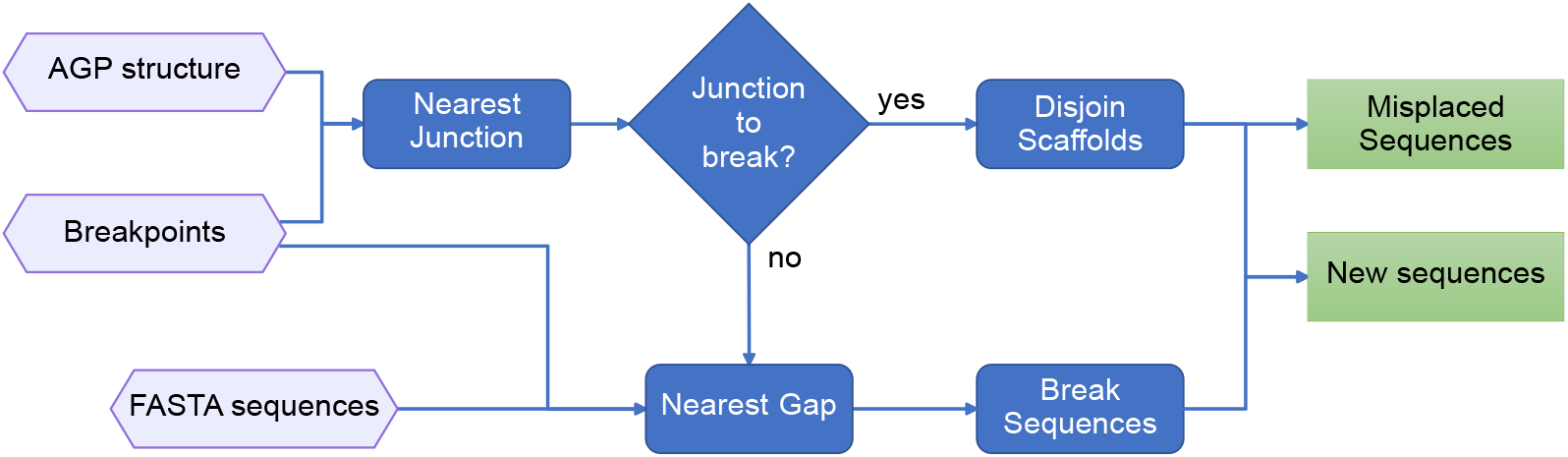
Schematic representation of HaploBreak procedure. HaploBreak selects appropriate breakpoints in mis-assembled regions by either breaking sequences at the closest scaffolding junction or gap. i) Given a pair of breakpoint coordinates, HaploBreak searches for the closest junction. If a scaffolding junction is found, it is associated with the breakpoint. Otherwise, the breakpoint is associated with the closest gap. ii) Then, HaploBreak breaks the assembly. If a pair of coordinates is associated with two different scaffolding junctions, the sequence between the coordinates is classified as misplaced. Otherwise, the sequence is broken at the gap.

**Supplemental figure 8:**
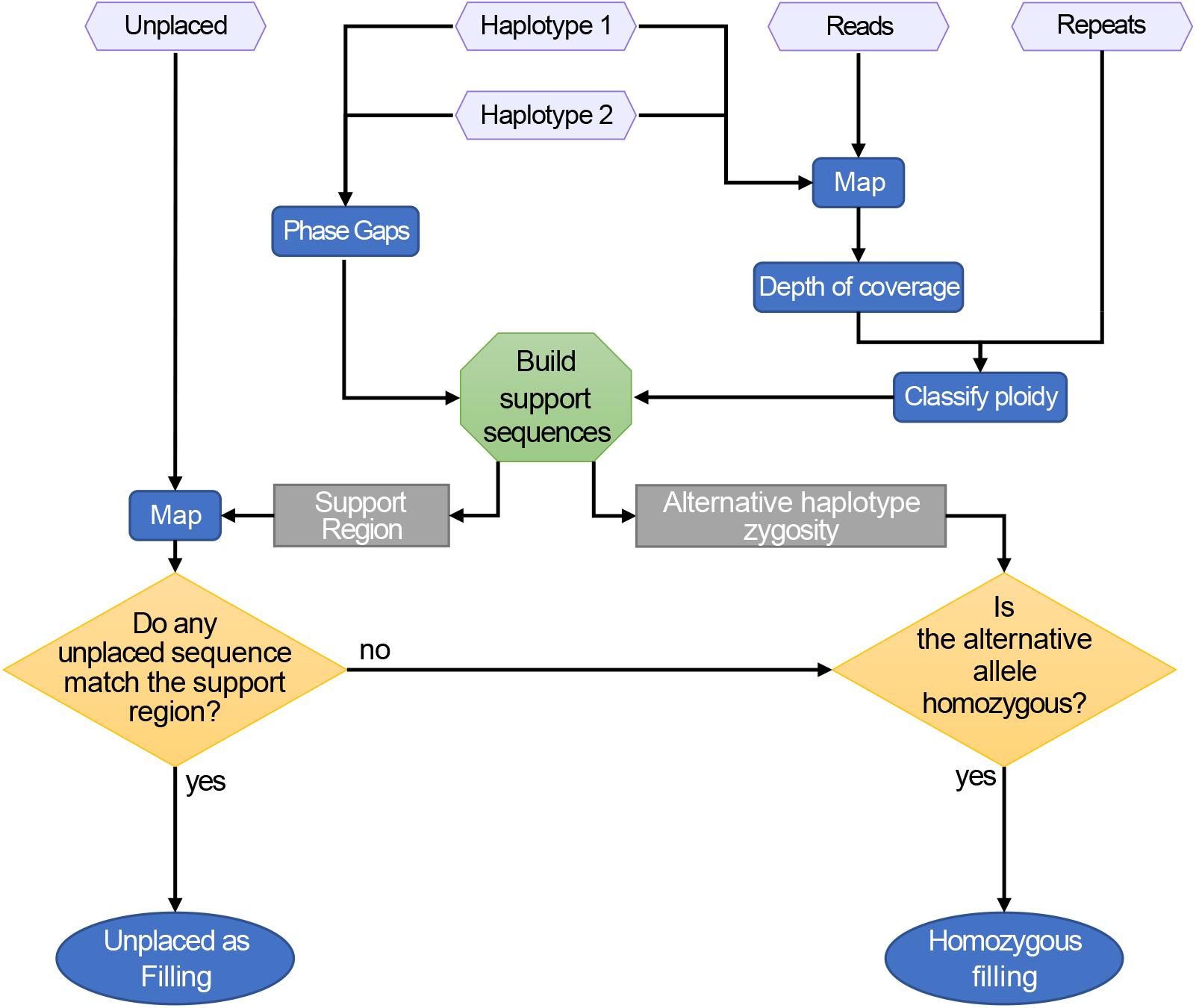
Schematic of the HaploFill procedure. HaploFill procedure: i) calculates sequencing coverage along each pseudomolecule, ii) evaluates the ploidy and repetitiveness along each pseudomolecule using the sequencing depth information and the repeat annotation, respectively, and iii) classifies each region of the genome based on the expected haploid depth of coverage; iv) for each gap, HaploFill identifies the corresponding region on the alternative haplotype and generates support sequences, and maps unplaced sequences on support sequences; v) the best filling information is assigned to the gap, prioritizing (in order of importance) the hybrid support region filler, the alternative support region filler, then the gap support region filler; vi) if the gap is diploid but cannot be filled, the region on the alternative haplotype corresponding to the gap is used as filler.

**Supplemental figure 9:**
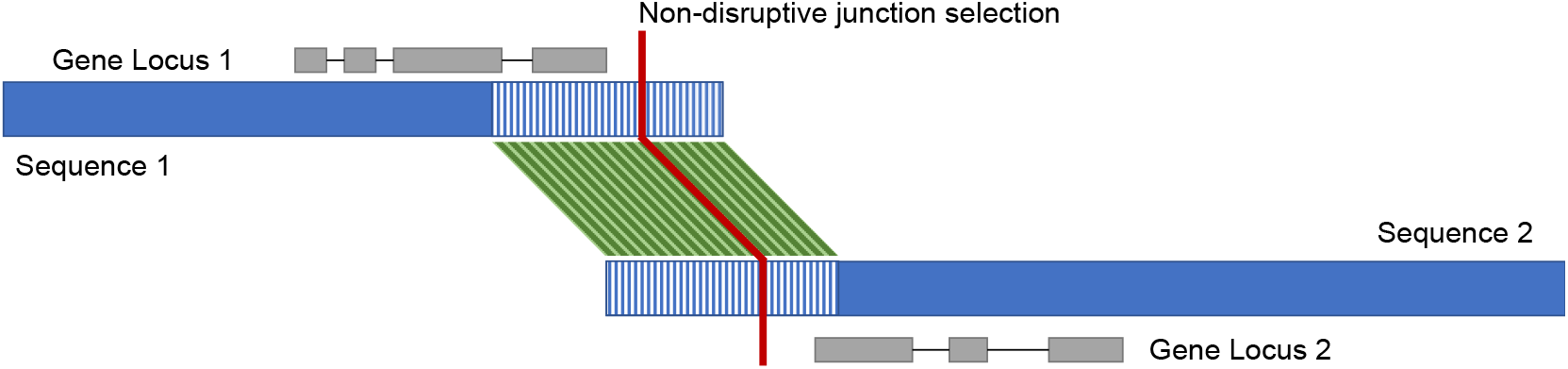
Schematic of the HaploMake procedure. HaploMake does an overlap-free sequence reconstruction without detrimentally affecting the gene annotation. When overlaps between sequences occur, the coordinates are corrected to avoid duplicating genomic content.

**Supplemental figure 10:**
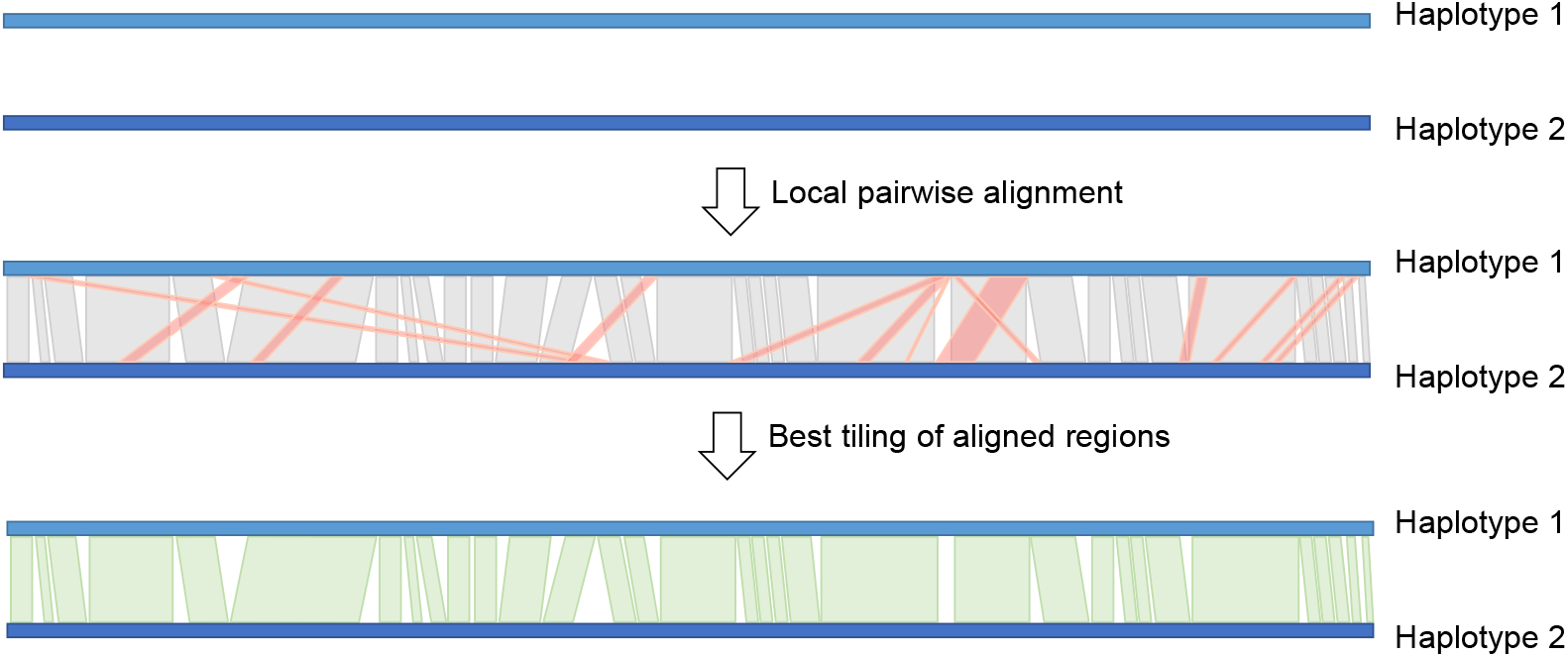
Schematic of the HaploMap procedure. HaploMap reports a map of the relationship between the two haplotypes. i) HaploMap performs local alignments between each pair of sequences, ii) the tiling path that maximizes the identity between both sequences is found, and iii) the coordinates of colinear regions in the bi-dimensional tiling path are reported.

**Supplemental figure 11:**
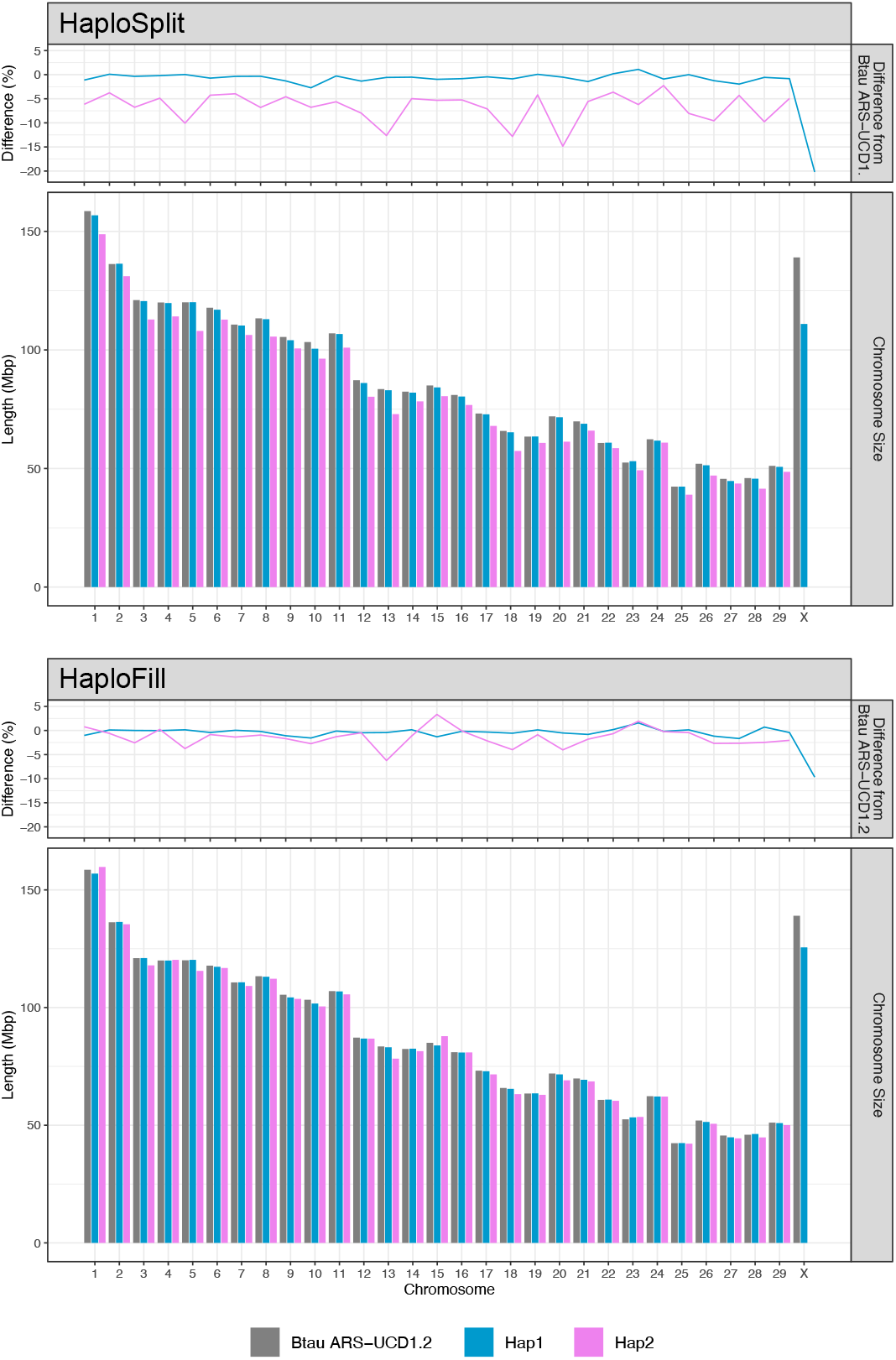
Results of B. taurus pseudomolecules reconstruction after HaploSplit and after HaploFill. Reconstructed pseudomolecule are compared in size to the ARS-UCD1.2 genome chromosome sequences. Each plots report a bar graph of the actual sizes and in terms of difference percentage from ARS-UCD1.2 genome chromosomes the expected genome size. In HaploSplit Haplotype 1 pseudomolecules deviate only by 0.7±0.7% form the expected size, Haplotype 2 by 6.7±3.0%. HaploFill further reduces the divergence, mostly in Haplotype 2 pseudomolecules where it goes down to 1.4±1.9%.

**Supplemental figure 12:**
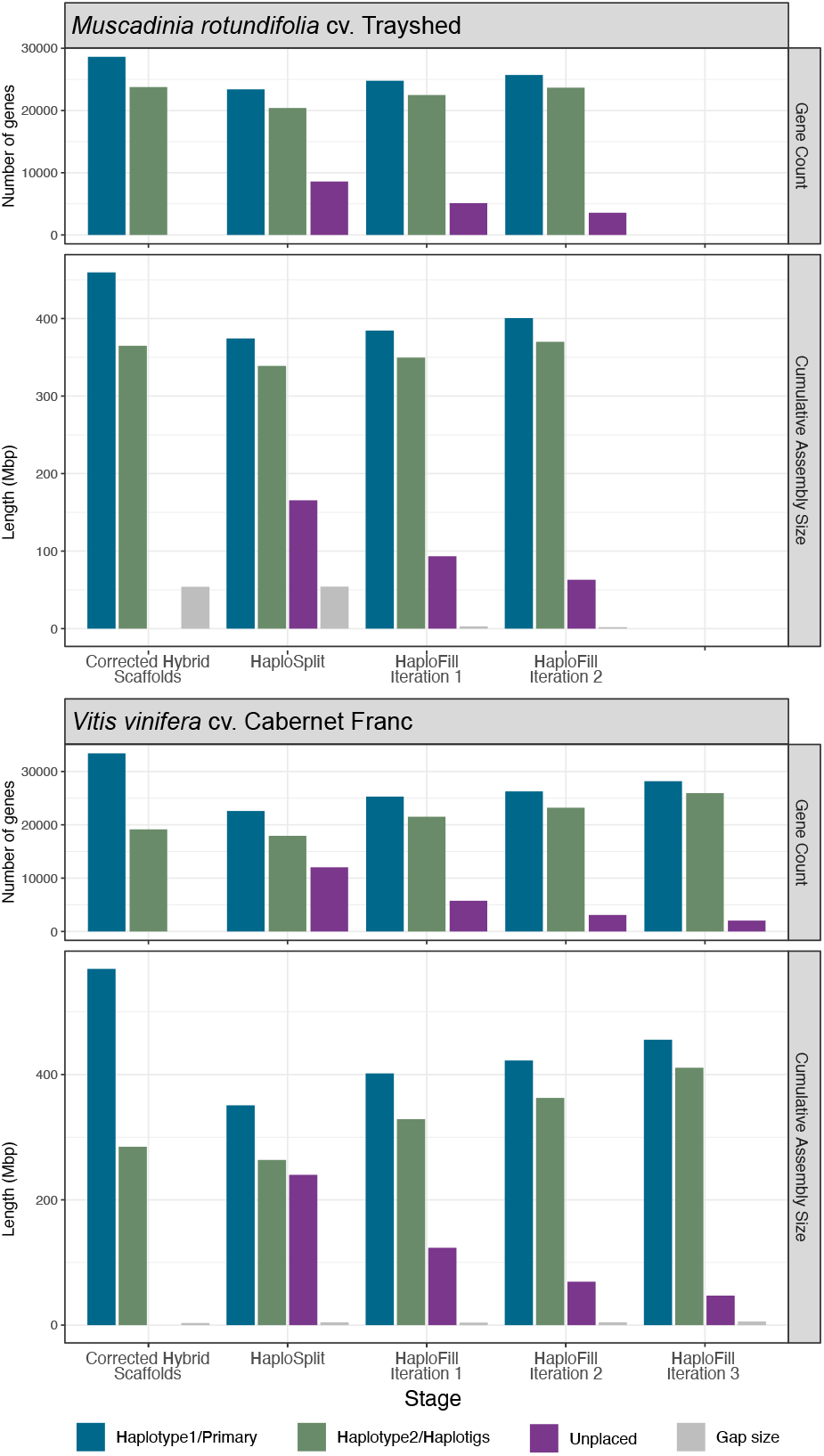
Results of HaploSync reconstruction of *M. rotundifolia* cv. Trayshed and *V. vinifera* cv. Cabernet Franc. Overview of pseudomolecule reconstruction results for the two Vitis species. The graphs reports both pseudomolecule sequence size and gene content at each step of HaploSync pipeline.

**Supplemental figure 13:**
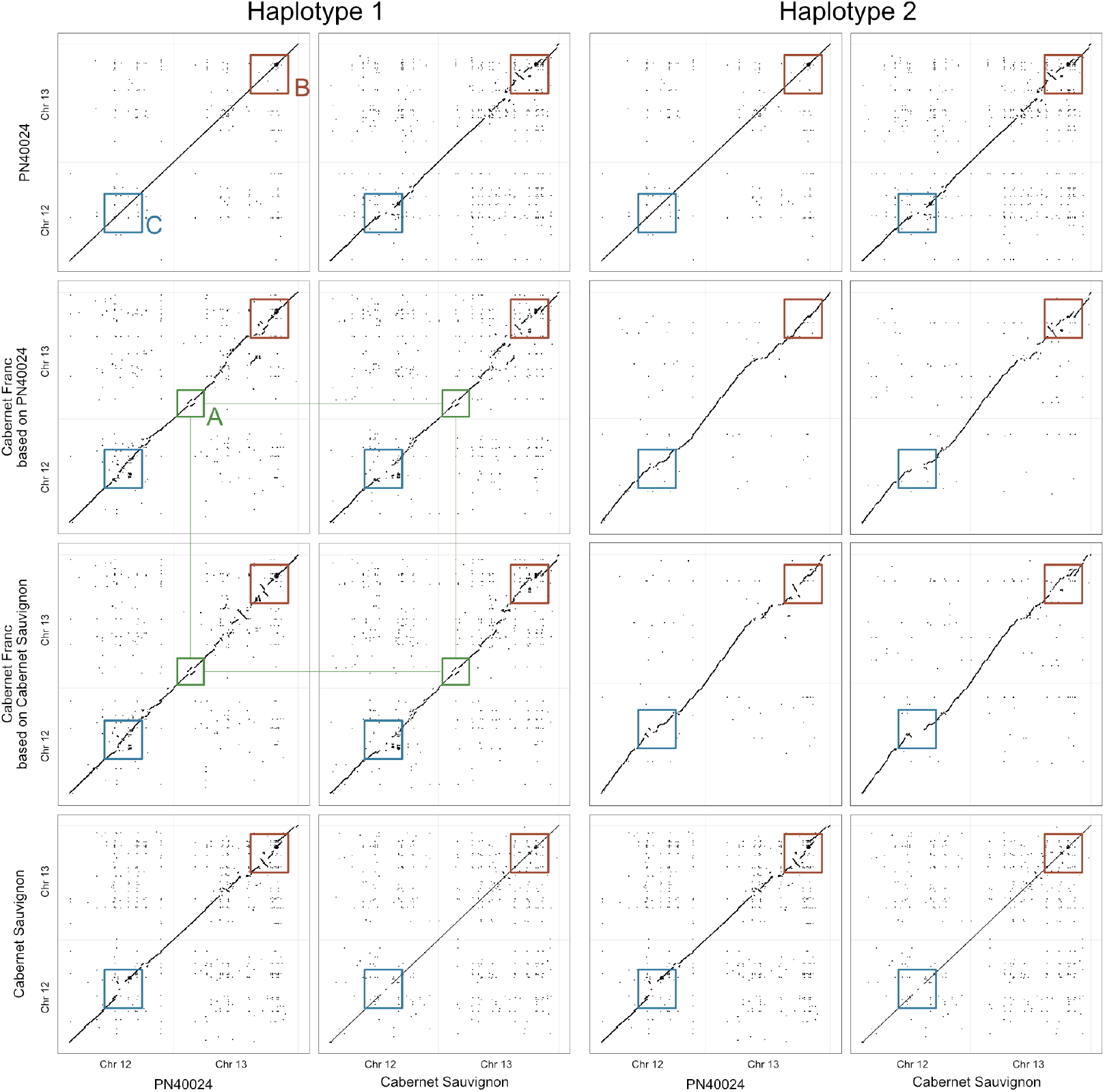
Guide genome overfitting. Dotplots comparing chromosome 12 and chromosome 13 of PN40024, Cabernet Sauvignon and Cabernet Franc, reconstructed with HaploSplit using PN40024 or Cabernet Sauvignon as guide genome, to the sequences of PN40024 and Cabernet Sauvignon. A boxes (green): Structural variants present in Cabernet Franc are reported in the results when part of long draft sequences. B boxes (red): Draft sequence location an orientation are placed accordance to the guide genome structure. With higher fragmentation (ex. sequences used for the second haplotype) increases also the overfit to the guide genome. C boxes (blue): Lack of information in the guide genome (gaps in Cabernet Sauvignon sequence) do not allow to insert place information inside the missing region unless the draft sequences do not anchor to the known part of the pseudomolecule.

**Supplemental figure 14:**
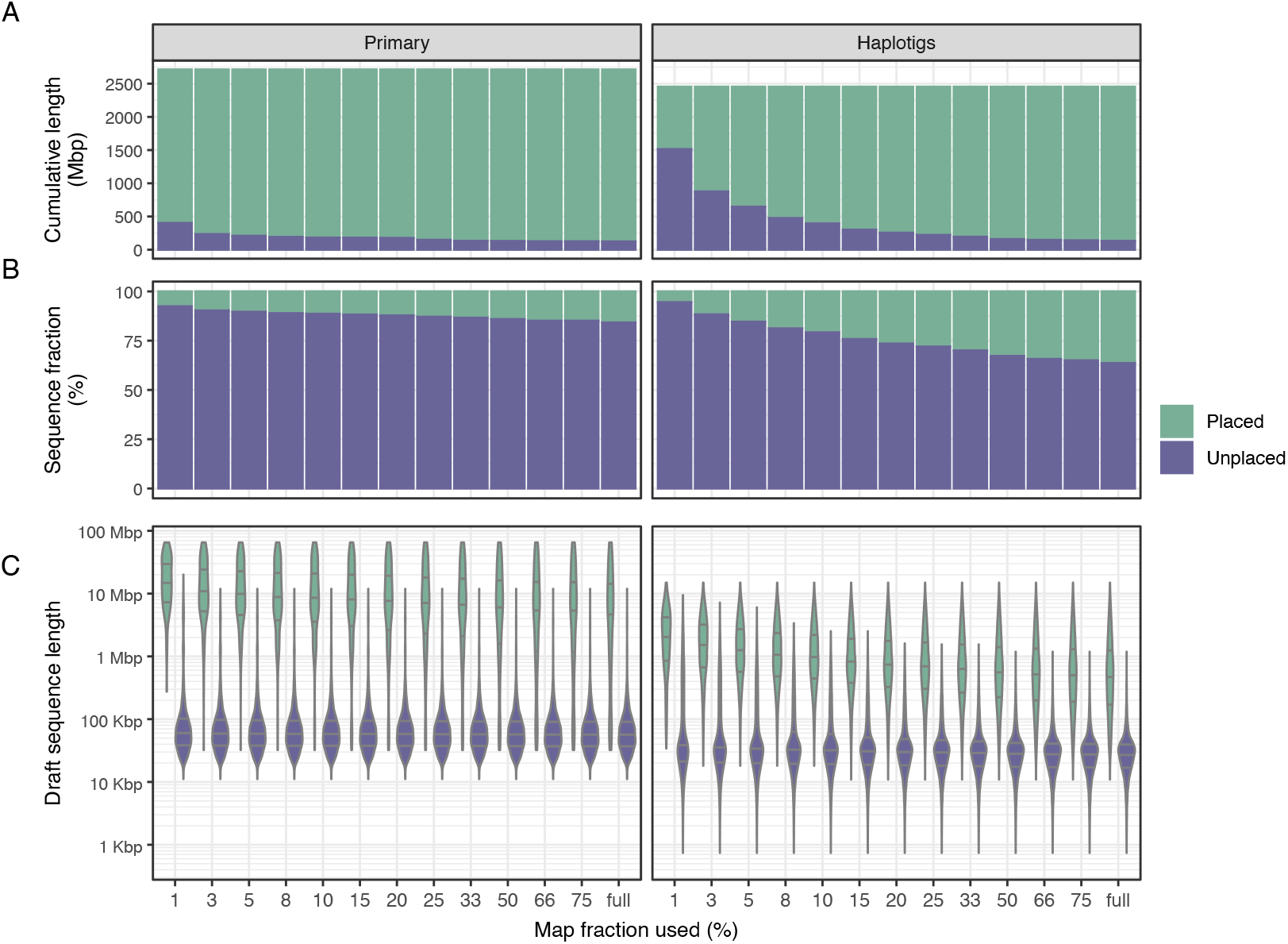
Performance of HaploSplit on markers subsampling. Overview of the HaploSplit performance in reconstructing *B. taurus* Angus x Brahma (12, 34) pseudomolecules using a subset of the full genetic map in terms of the input draft sequences placed. The results evidence the interrelated effect of map density and assembly fragmentation on the final assembly completeness, with only longer sequences used with the lowest amount of markers and shorter sequences requiring a higher density map to find a location. A) Cumulative length of placed and unplaced Primary sequences and Haplotigs with each marker dataset; B) Percentage of Primary sequences and Haplotigs finding a placement in the assembled pseudomolecules with each marker dataset; c) distribution of Primary sequences and Haplotigs lengths, placed and unplaced with each marker dataset.

**Supplemental table 1:**
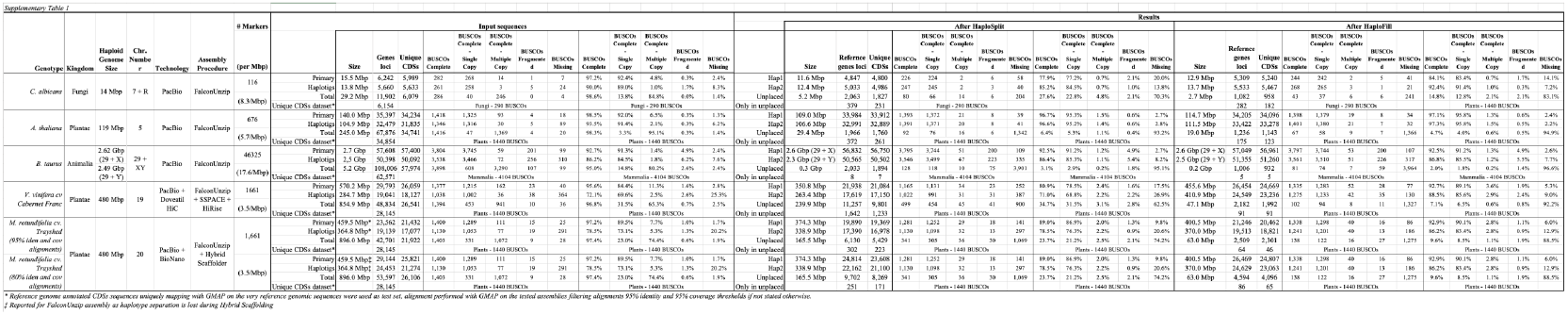
Gene content representation in the different assemblies. Table reporting the full statistics of the genomes assembled as testing dataset for HaploSync. C. albicans, A. thaliana and B. taurus genomes gene counts were obtained by mapping unique CDS sequences from the respective reference genome annotations (*C. albicans* SC5314_A22, *A. thaliana* TAIR10, *B. Taurus* Btau_ARS-UCD1.2) using GMAP (35). The unique gene mapping datasets were obtained by mapping the CDSs sequences of reference genome annotations to the respective reference genome, CDSs mapping in multiple locations in the haploid genome were removed form the dataset. Gene counts reported for *V. vinifera* cv Cabernet Franc and *M. rotundifolia* cv. Trayshed were obtained after performing the whole genome annotation as reported in (37) and (36) respectively.

**Supplemental table 2:**
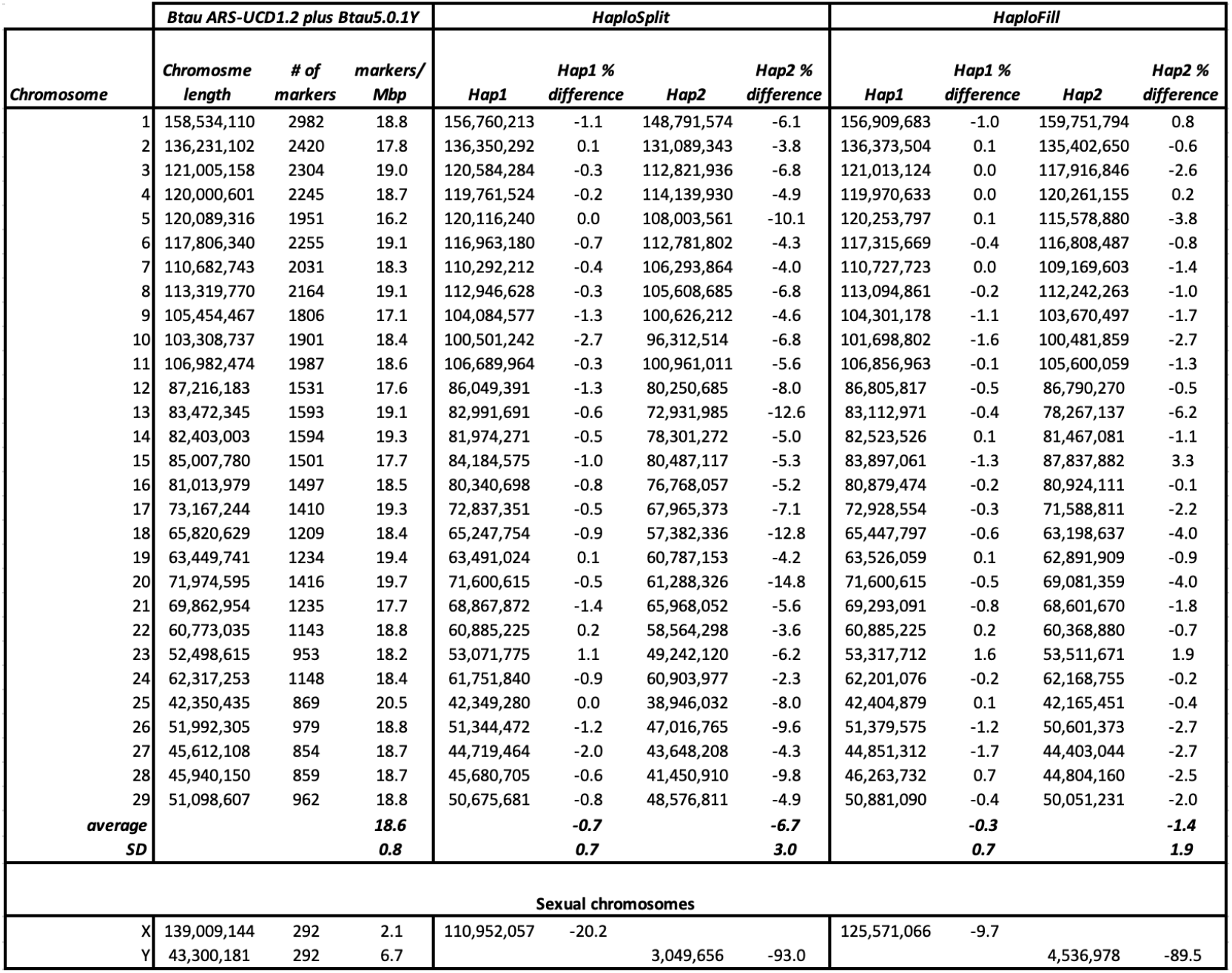
Results of B. taurus pseudomolecules reconstruction after HaploSplit and after HaploFill. Reconstructed pseudomolecule are compared in size to the ARS-UCD1.2 genome chromosome sequences. Each plots report a bar graph of the actual sizes and in terms of difference percentage from ARS-UCD1.2 genome chromosomes the expected genome size. In HaploSplit Haplotype 1 pseudomolecules deviate only by 0.7±0.7% form the expected size, Haplotype 2 by 6.7±3.0%. HaploFill further reduces the divergence, mostly in Haplotype 2 pseudomolecules where it goes down to 1.4±1.9%.

| Chromosome | Btau ARS-UCD1.2 plus Btau5.0.1Y |  |  | HaploSplit |  |  |  | HaploFill |  |  |  |
| --- | --- | --- | --- | --- | --- | --- | --- | --- | --- | --- | --- |
|  | Chromosome length | # of markers | markers/Mbp | Hap1 | Hap1 % difference | Hap2 | Hap2 % difference | Hap1 | Hap1 % difference | Hap2 | Hap2 % difference |
| 1 | 158,534,110 | 2982 | 18.8 | 156,760,213 | -1.1 | 148,791,574 | -6.1 | 156,909,683 | -1.0 | 159,751,794 | 0.8 |
| 2 | 136,231,102 | 2420 | 17.8 | 136,350,292 | 0.1 | 131,089,343 | -3.8 | 136,373,504 | 0.1 | 135,402,650 | -0.6 |
| 3 | 121,005,158 | 2304 | 19.0 | 120,584,284 | -0.3 | 112,821,936 | -6.8 | 121,013,124 | 0.0 | 117,916,846 | -2.6 |
| 4 | 120,000,601 | 2245 | 18.7 | 119,761,524 | -0.2 | 114,139,930 | -4.9 | 119,970,633 | 0.0 | 120,261,155 | 0.2 |
| 5 | 120,089,316 | 1951 | 16.2 | 120,116,240 | 0.0 | 108,003,561 | -10.1 | 120,253,797 | 0.1 | 115,578,880 | -3.8 |
| 6 | 117,806,340 | 2255 | 19.1 | 116,963,180 | -0.7 | 112,781,802 | -4.3 | 117,315,669 | -0.4 | 116,808,487 | -0.8 |
| 7 | 110,682,743 | 2031 | 18.3 | 110,292,212 | -0.4 | 106,293,864 | -4.0 | 110,727,723 | 0.0 | 109,169,603 | -1.4 |
| 8 | 113,319,770 | 2164 | 19.1 | 112,946,628 | -0.3 | 105,608,685 | -6.8 | 113,094,861 | -0.2 | 112,242,263 | -1.0 |
| 9 | 105,454,467 | 1806 | 17.1 | 104,084,577 | -1.3 | 100,626,212 | -4.6 | 104,301,178 | -1.1 | 103,670,497 | -1.7 |
| 10 | 103,308,737 | 1901 | 18.4 | 100,501,242 | -2.7 | 96,312,514 | -6.8 | 101,698,802 | -1.6 | 100,481,859 | -2.7 |
| 11 | 106,982,474 | 1987 | 18.6 | 106,689,964 | -0.3 | 100,961,011 | -5.6 | 106,856,963 | -0.1 | 105,600,059 | -1.3 |
| 12 | 87,216,183 | 1531 | 17.6 | 86,049,391 | -1.3 | 80,250,685 | -8.0 | 86,805,817 | -0.5 | 86,790,270 | -0.5 |
| 13 | 83,472,345 | 1593 | 19.1 | 82,991,691 | -0.6 | 72,931,985 | -12.6 | 83,112,971 | -0.4 | 78,267,137 | -6.2 |
| 14 | 82,403,003 | 1594 | 19.3 | 81,974,271 | -0.5 | 78,301,272 | -5.0 | 82,523,526 | 0.1 | 81,467,081 | -1.1 |
| 15 | 85,007,780 | 1501 | 17.7 | 84,184,575 | -1.0 | 80,487,117 | -5.3 | 83,897,061 | -1.3 | 87,837,882 | 3.3 |
| 16 | 81,013,979 | 1497 | 18.5 | 80,340,698 | -0.8 | 76,768,057 | -5.2 | 80,879,474 | -0.2 | 80,924,111 | -0.1 |
| 17 | 73,167,244 | 1410 | 19.3 | 72,837,351 | -0.5 | 67,965,373 | -7.1 | 72,928,554 | -0.3 | 71,588,811 | -2.2 |
| 18 | 65,820,629 | 1209 | 18.4 | 65,247,754 | -0.9 | 57,382,336 | -12.8 | 65,447,797 | -0.6 | 63,198,637 | -4.0 |
| 19 | 63,449,741 | 1234 | 19.4 | 63,491,024 | 0.1 | 60,787,153 | -4.2 | 63,526,059 | 0.1 | 62,891,909 | -0.9 |
| 20 | 71,974,595 | 1416 | 19.7 | 71,600,615 | -0.5 | 61,288,326 | -14.8 | 71,600,615 | -0.5 | 69,081,359 | -4.0 |
| 21 | 69,862,954 | 1235 | 17.7 | 68,867,872 | -1.4 | 65,968,052 | -5.6 | 69,293,091 | -0.8 | 68,601,670 | -1.8 |
| 22 | 60,773,035 | 1143 | 18.8 | 60,885,225 | 0.2 | 58,564,298 | -3.6 | 60,885,225 | 0.2 | 60,368,880 | -0.7 |
| 23 | 52,498,615 | 953 | 18.2 | 53,071,775 | 1.1 | 49,242,120 | -6.2 | 53,317,712 | 1.6 | 53,511,671 | 1.9 |
| 24 | 62,317,253 | 1148 | 18.4 | 61,751,840 | -0.9 | 60,903,977 | -2.3 | 62,201,076 | -0.2 | 62,168,755 | -0.2 |
| 25 | 42,350,435 | 869 | 20.5 | 42,349,280 | 0.0 | 38,946,032 | -8.0 | 42,404,879 | 0.1 | 42,165,451 | -0.4 |
| 26 | 51,992,305 | 979 | 18.8 | 51,344,472 | -1.2 | 47,016,765 | -9.6 | 51,379,575 | -1.2 | 50,601,373 | -2.7 |
| 27 | 45,612,108 | 854 | 18.7 | 44,719,464 | -2.0 | 43,648,208 | -4.3 | 44,851,312 | -1.7 | 44,403,044 | -2.7 |
| 28 | 45,940,150 | 859 | 18.7 | 45,680,705 | -0.6 | 41,450,910 | -9.8 | 46,263,732 | 0.7 | 44,804,160 | -2.5 |
| 29 | 51,098,607 | 962 | 18.8 | 50,675,681 | -0.8 | 48,576,811 | -4.9 | 50,881,090 | -0.4 | 50,051,231 | -2.0 |
| average |  |  | 18.6 |  | -0.7 |  | -6.7 |  | -0.3 |  | -1.4 |
| SD |  |  | 0.8 |  | 0.7 |  | 3.0 |  | 0.7 |  | 1.9 |
| Sexual chromosomes |  |  |  |  |  |  |  |  |  |  |  |
| X | 139,009,144 | 292 | 2.1 | 110,952,057 | -20.2 |  |  | 125,571,066 | -9.7 |  |  |
| Y | 43,300,181 | 292 | 6.7 |  |  | 3,049,656 | -93.0 |  |  | 4,536,978 | -89.5 |

**Supplemental table 3:**
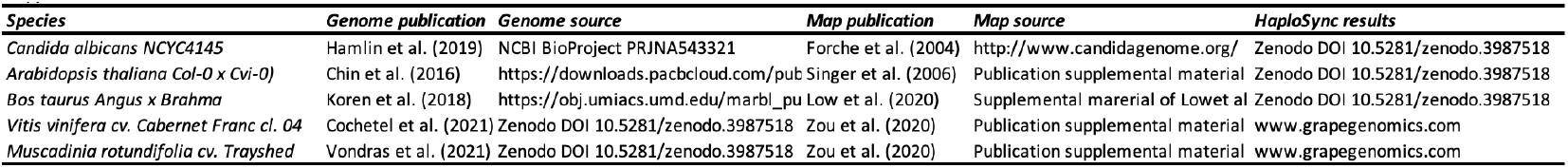
Data availability for input data and assembly results for testing datasets. Data availability for input genomic sequences, input maps used for each of testing datasets, and availability of the final assembly results.

## Bibliography

1. Anthony Rhoads and Kin Fai Au. PacBio Sequencing and its applications. Genomics, Proteomics and Bioinformatics, 13(5):278–289, 2015. ISSN 22103244. doi: 10.1016/j.gpb.2015.08.002.

2. Philipp Bongartz and Siegfried Schloissnig. Deep repeat resolution—the assembly of the Drosophila Histone Complex. Nucleic Acids Research, 47(3):e18–e18, feb 2019. ISSN 0305-1048. doi: 10.1093/nar/gky1194.

3. Huilong Du and Chengzhi Liang. Assembly of chromosome-scale contigs by efficiently resolving repetitive sequences with long reads. Nature Communications, 10(1):1–10, 2019. ISSN 20411723. doi: 10.1038/s41467-019-13355-3.

4. Pirita Paajanen, George Kettleborough, Elena López-Girona, Michael Giolai, Darren Heavens, David Baker, Ashleigh Lister, Fiorella Cugliandolo, Gail Wilde, Ingo Hein, Iain MacAulay, Glenn J. Bryan, and Matthew D. Clark. A critical comparison of technologies for a plant genome sequencing project. GigaScience, 8(3):1–10, 2019. ISSN 2047217X. doi: 10.1093/gigascience/giy163.

5. Kerrin S. Small, Michael Brudno, Matthew M. Hill, and Arend Sidow. A haplome alignment and reference sequence of the highly polymorphic ciona savignyi genome. Genome Biology, 8(3), 2007. ISSN 14747596. doi: 10.1186/gb-2007-8-3-r41.

6. Shengfeng Huang, Zelin Chen, Guangrui Huang, Ting Yu, Ping Yang, Jie Li, Yonggui Fu, Shaochun Yuan, Shangwu Chen, and Anlong Xu. HaploMerger: Reconstructing allelic relationships for polymorphic diploid genome assemblies. Genome Research, 22(8):1581–1588, 2012. ISSN 10889051. doi: 10.1101/gr.133652.111.

7. Alex Di Genova, Andrea M. Almeida, Claudia Muñoz-Espinoza, Paula Vizoso, Dante Travisany, Carol Moraga, Manuel Pinto, Patricio Hinrichsen, Ariel Orellana, and Alejandro Maass. Whole genome comparison between table and wine grapes reveals a comprehensive catalog of structural variants. BMC Plant Biology, 14(1), 2014. ISSN 14712229. doi: 10.1186/1471-2229-14-7.

8. Hideki Hirakawa, Kenta Shirasawa, Shunichi Kosugi, Kosuke Tashiro, Shinobu Nakayama, Manabu Yamada, Mistuyo Kohara, Akiko Watanabe, Yoshie Kishida, Tsunakazu Fujishiro, Hisano Tsuruoka, Chiharu Minami, Shigemi Sasamoto, Midori Kato, Keiko Nanri, Akiko Komaki, Tomohiro Yanagi, Qin Guoxin, Fumi Maeda, Masami Ishikawa, Satoru Kuhara, Shusei Sato, Satoshi Tabata, and Sachiko N. Isobe. Dissection of the octoploid strawberry genome by deep sequencing of the genomes of fragaria species. DNA Research, 21(2): 169–181, 2014. ISSN 17561663. doi: 10.1093/dnares/dst049.

9. Rei Kajitani, Kouta Toshimoto, Hideki Noguchi, Atsushi Toyoda, Yoshitoshi Ogura, Miki Okuno, Mitsuru Yabana, Masayuki Harada, Eiji Nagayasu, Haruhiko Maruyama, Yuji Kohara, Asao Fujiyama, Tetsuya Hayashi, and Takehiko Itoh. Efficient de novo assembly of highly heterozygous genomes from whole-genome shotgun short reads. Genome Research, 24 (8):1384–1395, 2014. ISSN 15495469. doi: 10.1101/gr.170720.113.

10. Hua Ying, David C. Hayward, Ira Cooke, Weiwen Wang, Aurelie Moya, Kirby R. Siemering, Susanne Sprungala, Eldon E. Ball, Sylvain Forêt, and David J. Miller. The whole-genome sequence of the coral acropora millepora. Genome Biology and Evolution, 11(5):1374–1379, 2019. ISSN 17596653. doi: 10.1093/gbe/evz077.

11. Shilpa Garg, Arkarachai Fungtammasan, Andrew Carroll, Mike Chou, Anthony Schmitt, Xiang Zhou, Stephen Mac, Paul Peluso, Emily Hatas, Jay Ghurye, Jared Maguire, Medhat Mahmoud, Haoyu Cheng, David Heller, Justin M. Zook, Tobias Moemke, Tobias Marschall, Fritz J. Sedlazeck, John Aach, Chen Shan Chin, George M. Church, and Heng Li. Chromosome-scale, haplotype-resolved assembly of human genomes. Nature Biotechnology, 39(3):309–312, 2021. ISSN 15461696. doi: 10.1038/s41587-020-0711-0.

12. Wai Yee Low, Rick Tearle, Ruijie Liu, Sergey Koren, Arang Rhie, Derek M. Bickhart, Benjamin D. Rosen, Zev N. Kronenberg, Sarah B. Kingan, Elizabeth Tseng, Françoise Thibaud-Nissen, Fergal J. Martin, Konstantinos Billis, Jay Ghurye, Alex R. Hastie, Joyce Lee, Andy W. C. Pang, Michael P. Heaton, Adam M. Phillippy, Stefan Hiendleder, Timothy P. L. Smith, and John L. Williams. Haplotype-resolved genomes provide insights into structural variation and gene content in Angus and Brahman cattle. Nature Communications, 11(1):2071, dec 2020. ISSN 2041-1723. doi: 10.1038/s41467-020-15848-y.

13. Mélanie Massonnet, Noé Cochetel, Andrea Minio, Amanda M. Vondras, Jerry Lin, Aline Muyle, Jadran F. Garcia, Yongfeng Zhou, Massimo Delledonne, Summaira Riaz, Rosa Figueroa-Balderas, Brandon S. Gaut, and Dario Cantu. The genetic basis of sex determination in grapes. Nature communications, 11(1):2902, dec 2020. ISSN 2041-1723. doi: 10.1038/s41467-020-16700-z.

14. Xuepeng Sun, Chen Jiao, Heidi Schwaninger, C. Thomas Chao, Yumin Ma, Naibin Duan, Awais Khan, Seunghyun Ban, Kenong Xu, Lailiang Cheng, and et al. Phased diploid genome assemblies and pan-genomes provide insights into the genetic history of apple domestication. Nature Genetics, 52(12):1423–1432, 2020. ISSN 15461718. doi: 10.1038/s41588-020-00723-9.

15. Peng Zhou, Zhi Li, Erika Magnusson, Fabio Gomez Cano, Peter A. Crisp, Jaclyn M. Noshay, Erich Grotewold, Candice N. Hirsch, Steven P. Briggs, and Nathan M. Springer. Meta gene regulatory networks in maize highlight functionally relevant regulatory interactions. Plant Cell, 32(5):1377–1396, 2020. ISSN 1532298X. doi: 10.1105/tpc.20.00080.

16. Ben N. Mansfeld, Adam Boyher, Jeffrey C. Berry, Mark Wilson, Shujun Ou, Seth Polydore, Todd P. Michael, Noah Fahlgren, and Rebecca S. Bart. Large structural variations in the haplotype-resolved african cassava genome. Plant Journal, 108(6):1830–1848, 2021. ISSN 1365313X. doi: 10.1111/tpj.15543.

17. Chen Shan Chin, Paul Peluso, Fritz J. Sedlazeck, Maria Nattestad, Gregory T. Concepcion, Alicia Clum, Christopher Dunn, Ronan O’Malley, Rosa Figueroa-Balderas, Abraham Morales-Cruz, Grant R. Cramer, Massimo Delledonne, Chongyuan Luo, Joseph R. Ecker, Dario Cantu, David R. Rank, and Michael C. Schatz. Phased diploid genome assembly with single-molecule real-time sequencing. Nature Methods, 13(12):1050–1054, 2016. ISSN 15487105. doi: 10.1038/nmeth.4035.

18. Andrea Minio, Mélanie Massonnet, Rosa Figueroa-Balderas, Alvaro Castro, and Dario Cantu. Diploid genome assembly of the wine grape carménère. G3: Genes, Genomes, Genetics, 9(5):1331–1337, may 2019. ISSN 21601836. doi: 10.1534/g3.119.400030.

19. Lorenzo Barchi, Marco Pietrella, Luca Venturini, Andrea Minio, Laura Toppino, Alberto Acquadro, Giuseppe Andolfo, Giuseppe Aprea, Carla Avanzato, Laura Bassolino, Cinzia Comino, Alessandra Dal Molin, Alberto Ferrarini, Louise Chappell Maor, Ezio Portis, Sebastian Reyes-Chin-Wo, Riccardo Rinaldi, Tea Sala, Davide Scaglione, Prashant Sonawane, Paola Tononi, Efrat Almekias-Siegl, Elisa Zago, Maria Raffaella Ercolano, Asaph Aharoni, Massimo Delledonne, Giovanni Giuliano, Sergio Lanteri, and Giuseppe Leonardo Rotino. A chromosome-anchored eggplant genome sequence reveals key events in Solanaceae evolution. Scientific Reports, 9(1):1–13, 2019. ISSN 20452322. doi: 10.1038/s41598-019-47985-w.

20. Prashant S Hosmani, Mirella Flores-Gonzalez, Henri van de Geest, Florian Maumus, Linda V Bakker, Elio Schijlen, Jan van Haarst, Jan Cordewener, Gabino Sanchez-Perez, Sander Peters, and et al. An improved de novo assembly and annotation of the tomato reference genome using single-molecule sequencing, hi-c proximity ligation and optical maps. bioRxiv, 2012:767764, 2019. doi: 10.1101/767764.

21. Andreas Wallberg, Ignas Bunikis, Olga Vinnere Pettersson, Mai-Britt Mosbech, Anna K. Childers, Jay D. Evans, Alexander S. Mikheyev, Hugh M. Robertson, Gene E. Robinson, and Matthew T. Webster. A hybrid de novo genome assembly of the honeybee, Apis mellifera, with chromosome-length scaffolds. BMC Genomics, 20(1):275, dec 2019. ISSN 1471-2164. doi: 10.1186/s12864-019-5642-0.

22. Karen H. Miga, Sergey Koren, Arang Rhie, Mitchell R. Vollger, Ariel Gershman, Andrey Bzikadze, Shelise Brooks, Edmund Howe, David Porubsky, Glennis A. Logsdon, Valerie A. Schneider, Tamara Potapova, Jonathan Wood, William Chow, Joel Armstrong, Jeanne Fredrickson, Evgenia Pak, Kristof Tigyi, Milinn Kremitzki, Christopher Markovic, Valerie Maduro, Amalia Dutra, Gerard G. Bouffard, Alexander M. Chang, Nancy F. Hansen, Amy B. Wilfert, Françoise Thibaud-Nissen, Anthony D. Schmitt, Jon-Matthew Belton, Siddarth Selvaraj, Megan Y. Dennis, Daniela C. Soto, Ruta Sahasrabudhe, Gulhan Kaya, Josh Quick, Nicholas J. Loman, Nadine Holmes, Matthew Loose, Urvashi Surti, Rosa ana Risques, Tina A. Graves Lindsay, Robert Fulton, Ira Hall, Benedict Paten, Kerstin Howe, Winston Timp, Alice Young, James C. Mullikin, Pavel A. Pevzner, Jennifer L. Gerton, Beth A. Sullivan, Evan E. Eichler, and Adam M. Phillippy. Telomere-to-telomere assembly of a complete human X chromosome. Nature, 585(7823):79–84, sep 2020. ISSN 0028-0836. doi: 10.1038/s41586-020-2547-7.

23. Jaebum Kim, Denis M. Larkin, Qingle Cai, Asan, Yongfen Zhang, Ri Li Ge, Loretta Auvil, Boris Capitanu, Guojie Zhang, Harris A. Lewin, and Jian Ma. Reference-assisted chromosome assembly. Proceedings of the National Academy of Sciences of the United States of America, 110(5):1785–1790, 2013. ISSN 00278424. doi: 10.1073/pnas.1220349110.

24. Haibao Tang, Xingtan Zhang, Chenyong Miao, Jisen Zhang, Ray Ming, James C. Schnable, Patrick S. Schnable, Eric Lyons, and Jianguo Lu. ALLMAPS: Robust scaffold ordering based on multiple maps. Genome Biology, 16(1):1–15, 2015. ISSN 1474760X. doi: 10.1186/s13059-014-0573-1.

25. Gaik Tamazian, Pavel Dobrynin, Ksenia Krasheninnikova, Aleksey Komissarov, Klaus Peter Koepfli, and Stephen J. O’Brien. Chromosomer: A reference-based genome arrangement tool for producing draft chromosome sequences. GigaScience, 5(1):1–11, 2016. ISSN 2047217X. doi: 10.1186/s13742-016-0141-6.

26. Michael Alonge, Sebastian Soyk, Srividya Ramakrishnan, Xingang Wang, Sara Goodwin, Fritz J. Sedlazeck, Zachary B. Lippman, and Michael C. Schatz. RaGOO: Fast and accurate reference-guided scaffolding of draft genomes. Genome Biology, 20(1):1–17, 2019. ISSN 1474760X. doi: 10.1186/s13059-019-1829-6.

27. Yi Ren, Hong Zhao, Qinghe Kou, Jiao Jiang, Shaogui Guo, Haiying Zhang, Wenju Hou, Xiaohua Zou, Honghe Sun, Guoyi Gong, Amnon Levi, and Yong Xu. A high resolution genetic map anchoring scaffolds of the sequenced watermelon genome. PLoS ONE, 7(1), 2012. ISSN 19326203. doi: 10.1371/journal.pone.0029453.

28. Heng Li. Minimap2: Pairwise alignment for nucleotide sequences. Bioinformatics, 34(18): 3094–3100, sep 2018. ISSN 14602059. doi: 10.1093/bioinformatics/bty191.

29. Aaron R. Quinlan and Ira M. Hall. BEDTools: A flexible suite of utilities for comparing genomic features. Bioinformatics, 26(6):841–842, mar 2010. ISSN 13674803. doi: 10.1093/bioinformatics/btq033.

30. Guillaume Marçais, Arthur L. Delcher, Adam M. Phillippy, Rachel Coston, Steven L. Salzberg, and Aleksey Zimin. MUMmer4: A fast and versatile genome alignment system. PLoS Computational Biology, 14(1):e1005944, jan 2018. ISSN 15537358. doi: 10.1371/journal.pcbi.1005944.

31. Jennafer A.P. Hamlin, Guilherme B. Dias, Casey M. Bergman, and Douda Bensasson. Phased diploid genome assemblies for three strains of Candida albicans from oak trees. G3: Genes, Genomes, Genetics, 9(11):3547–3554, 2019. ISSN 21601836. doi: 10.1534/g3.119.400486.

32. Anja Forche, P T Magee, B. B. Magee, and Georgiana May. Genome-wide single-nucleotide polymorphism map for Candida albicans. Eukaryotic cell, 3(3):705–14, jun 2004. ISSN 1535-9778. doi: 10.1128/EC.3.3.705-714.2004.

33. Tatjana Singer, Yiping Fan, Hur Song Chang, Tong Zhu, Samuel P. Hazen, and Steven P. Briggs. A high-resolution map of Arabidopsis recombinant inbred lines by whole-genome exon array hybridization. PLoS Genetics, 2(9):1352–1361, 2006. ISSN 15537390. doi: 10.1371/journal.pgen.0020144.

34. Sergey Koren, Arang Rhie, Brian P. Walenz, Alexander T. Dilthey, Derek M. Bickhart, Sarah B. Kingan, Stefan Hiendleder, John L. Williams, Timothy P L Smith, and Adam M. Phillippy. De novo assembly of haplotype-resolved genomes with trio binning. Nature Biotechnology, 36(12):1174–1182, dec 2018. ISSN 1087-0156. doi: 10.1038/nbt.4277.

35. Thomas D. Wu and Colin K. Watanabe. GMAP: A genomic mapping and alignment program for mRNA and EST sequences. Bioinformatics, 21(9):1859–1875, 2005. ISSN 13674803. doi: 10.1093/bioinformatics/bti310.

36. Amanda M. Vondras, Larry Lerno, Mélanie Massonnet, Andrea Minio, Adib Rowhani, Dingren Liang, Jadran Garcia, Daniela Quiroz, Rosa Figueroa-Balderas, Deborah A. Golino, Susan E. Ebeler, Maher Al Rwahnih, and Dario Cantu. Rootstock influences the effect of grapevine leafroll-associated viruses on berry development and metabolism via abscisic acid signalling. Molecular Plant Pathology, 22(8):984–1005, 2021. ISSN 13643703. doi: 10.1111/mpp.13077.

37. Noé Cochetel, Andrea Minio, Mélanie Massonnet, Amanda M Vondras, Rosa Figueroa-Balderas, and Dario Cantu. Diploid chromosome-scale assembly of the Muscadinia rotundifolia genome supports chromosome fusion and disease resistance gene expansion during Vitis and Muscadinia divergence. G3 Genes|Genomes|Genetics, 11(4), apr 2021. ISSN 2160-1836. doi: 10.1093/g3journal/jkab033.

38. Cheng Zou, Avinash Karn, Bruce Reisch, Allen Nguyen, Yongming Sun, Yun Bao, Michael S. Campbell, Deanna Church, Stephen Williams, Xia Xu, Craig A. Ledbetter, Sagar Patel, Anne Fennell, Jeffrey C. Glaubitz, Matthew Clark, Doreen Ware, Jason P. Londo, Qi Sun, and Lance Cadle-Davidson. Haplotyping the Vitis collinear core genome with rhAmpSeq improves marker transferability in a diverse genus. Nature Communications, 11(1):413, dec 2020. ISSN 2041-1723. doi: 10.1038/s41467-019-14280-1.

39. Benjamin D. Rosen, Derek M. Bickhart, Robert D. Schnabel, Sergey Koren, Christine G. Elsik, Elizabeth Tseng, Troy N. Rowan, Wai Y. Low, Aleksey Zimin, Christine Couldrey, Richard Hall, Wenli Li, Arang Rhie, Jay Ghurye, Stephanie D. McKay, Françoise Thibaud-Nissen, Jinna Hoffman, Brenda M. Murdoch, Warren M. Snelling, Tara G. McDaneld, John A. Hammond, John C. Schwartz, Wilson Nandolo, Darren E. Hagen, Christian Dreischer, Sebastian J. Schultheiss, Steven G. Schroeder, Adam M. Phillippy, John B. Cole, Curtis P Van Tassell, George Liu, Timothy P L Smith, and Juan F. Medrano. De novo assembly of the cattle reference genome with single-molecule sequencing. GigaScience, 9(3):1–9, mar 2020. ISSN 2047-217X. doi: 10.1093/gigascience/giaa021.

40. Riccardo Velasco, Andrey Zharkikh, Michela Troggio, Dustin A. Cartwright, Alessandro Cestaro, Dmitry Pruss, Massimo Pindo, Lisa M Fitzgerald, Silvia Vezzulli, Julia Reid, Giulia Malacarne, Diana Iliev, Giuseppina Coppola, Bryan Wardell, Diego Micheletti, Teresita Macalma, Marco Facci, Jeff T. Mitchell, Michele Perazzolli, Glenn Eldredge, Pamela Gatto, Rozan Oyzerski, Marco Moretto, Natalia Gutin, Marco Stefanini, Yang Chen, Cinzia Segala, Christine Davenport, Lorenzo Demattè, Amy Mraz, Juri Battilana, Keith Stormo, Fabrizio Costa, Quanzhou Tao, Azeddine Si-Ammour, Tim Harkins, Angie Lackey, Clotilde Perbost, Bruce Taillon, Alessandra Stella, Victor Solovyev, Jeffrey A. Fawcett, Lieven Sterck, Klaas Vandepoele, Stella M. Grando, Stefano Toppo, Claudio Moser, Jerry Lanchbury, Robert Bogden, Mark Skolnick, Vittorio Sgaramella, Satish K. Bhatnagar, Paolo Fontana, Alexander Gutin, Yves Van de Peer, Francesco Salamini, and Roberto Viola. A high quality draft consensus sequence of the genome of a heterozygous grapevine variety. PloS one, 2(12): e1326, dec 2007. ISSN 1932-6203. doi: 10.1371/journal.pone.0001326.

41. Michael J. Roach, Simon A. Schmidt, and Anthony R. Borneman. Purge Haplotigs: allelic contig reassignment for third-gen diploid genome assemblies. BMC Bioinformatics, 19(1): 460, dec 2018. ISSN 1471-2105. doi: 10.1186/s12859-018-2485-7.

42. Amanda M. Vondras, Andrea Minio, Barbara Blanco-Ulate, Rosa Figueroa-Balderas, Michael A. Penn, Yongfeng Zhou, Danelle Seymour, Zirou Ye, Dingren Liang, Lucero K. Espinoza, Michael M. Anderson, M. Andrew Walker, Brandon Gaut, and Dario Cantu. The genomic diversification of grapevine clones. BMC genomics, 20(1):972, 2019. ISSN 14712164. doi: 10.1186/s12864-019-6211-2.

43. Yongfeng Zhou, Andrea Minio, Mélanie Massonnet, Edwin Solares, Yuanda Lv, Tengiz Beridze, Dario Cantu, and Brandon S. Gaut. The population genetics of structural variants in grapevine domestication. Nature Plants, 5(9):965–979, 2019. ISSN 20550278. doi: 10.1038/s41477-019-0507-8.

44. Olivier Jaillon, Jean Marc Aury, Benjamin Noel, Alberto Policriti, Christian Clepet, Alberto Casagrande, Nathalie Choisne, Sébastien Aubourg, Nicola Vitulo, Claire Jubin, Alessandro Vezzi, Fabrice Legeai, Philippe Hugueney, Corinne Dasilva, David Horner, Erica Mica, Delphine Jublot, Julie Poulain, Clémence Bruyère, Alain Billault, Béatrice Segurens, Michel Gouyvenoux, Edgardo Ugarte, Federica Cattonaro, Véronique Anthouard, Virginie Vico, Cristian Del Fabbro, Michaël Alaux, Gabriele Di Gaspero, Vincent Dumas, Nicoletta Felice, Sophie Paillard, Irena Juman, Marco Moroldo, Simone Scalabrin, Aurélie Canaguier, Isabelle Le Clainche, Giorgio Malacrida, Eléonore Durand, Graziano Pesole, Valérie Laucou, Philippe Chatelet, Didier Merdinoglu, Massimo Delledonne, Mario Pezzotti, Alain Lecharny, Claude Scarpelli, François Artiguenave, M. Enrico Pè, Giorgio Valle, Michele Morgante, Michel Caboche, Anne Françoise Adam-Blondon, Jean Weissenbach, Francis Quétier, and Patrick Wincker. The grapevine genome sequence suggests ancestral hexaploidization in major angiosperm phyla. Nature, 449(7161):463–467, sep 2007. ISSN 14764687. doi: 10.1038/nature06148.

45. Aurélie Canaguier, Jérôme Grimplet, Gabriele Di Gaspero, Simone Scalabrin, Éric Duchêne, Nathalie Choisne, Nacer Mohellibi, Cécile Guichard, Stéphane Rombauts, Isabelle Le Clainche, Aurélie Bérard, Aurélie Chauveau, Rémi Bounon, Camille Rustenholz, Michele Morgante, Marie Christine Le Paslier, Dominique Brunel, and Anne-Françoise Adam-Blondon. A new version of the grapevine reference genome assembly (12X.v2) and of its annotation (VCost.v3). Genomics data, 14(July):56–62, dec 2017. ISSN 2213-5960. doi: 10.1016/j.gdata.2017.09.002.

46. Sergey Koren, Arang Rhie, Brian P. Walenz, Alexander T. Dilthey, Derek M. Bickhart, Sarah B. Kingan, Stefan Hiendleder, John L. Williams, Timothy P.L. Smith, and Adam M. Phillippy. De novo assembly of haplotype-resolved genomes with trio binning. Nature Biotechnology, 2018. ISSN 15461696. doi: 10.1038/nbt.4277.

